# Genomic insights into the evolutionary history and diversification of water voles (*Arvicola* spp.)

**DOI:** 10.64898/2026.11.25.769000

**Authors:** Aleksandra Deroń, Amanda Lindahl, Barbara Bujalska, Ahmad Mahmoudi, Boris Krystufek, Armando Nappi, Aleksander Bukhnikashvili, Soraia Barbosa, Damien Flas, Monika Knul, Boyan Milchev, Karl Frafjord, Miriam Belmaker, Jan Decher, Daniela C. Kalthoff, John Stewart, Jan M Wójcik, Emmanuel Desclaux, Claudio Berto, Adam Nadachowski, Love Dalén, Mateusz Baca

## Abstract

Water voles (*Arvicola* spp.) exhibit substantial morphological, ecological, and genetic variation which has initiated a longstanding debate over their evolutionary history and taxonomic status. Although five species have recently been proposed; *A. amphibius*, *A. sapidus*, *A. persicus*, *A. italicus* and *A. monticola*, their evolutionary relationships and divergence times remain poorly resolved. Here, we analyse 115 complete mitochondrial genomes and nuclear genomic data from 45 individuals representing all currently proposed species. We estimate that modern water voles began diverging ca. 400 ka ago, considerably more recently than previously estimated but broadly consistent with the fossil record. Further diversification occurred during the Middle and Late Pleistocene. However, we detect multiple discordances between the mitochondrial and nuclear relationships, and admixture analyses suggest that there has been historical gene flow between lineages. Furthermore, we find lower long-term effective population sizes and reduced genetic variation in several southern populations, potentially reflecting more restricted habitat availability or greater isolation than in central and northern populations. Additionally, in Georgia we identify a sister lineage to *A. persicus* and show that Anatolian water voles actually belong to this lineage and not *A. amphibius*. Our findings offer new insights into the complex evolutionary history of water voles, which was initially shaped by Pleistocene climatic fluctuations and subsequently reshaped by gene flow.

## 2 Introduction

Rodents are the most diverse group of mammals, comprising more than 40% of all mammalian species (Kay and Hoekstra, 2008). One reason for this diversity is the rapid genomic evolution of rodents, which for reference is about ∼4–6 times faster than primates (Churakov et al., 2010; Wu and Li, 1985). Their ability to evolve abundant interspecific phenotypic variation in response to different environments have made it possible for rodents to inhabit almost all continents and habitats on Earth (Kay and Hoekstra, 2008). The amount of variation makes rodent phylogenies difficult to resolve for many taxa, but also provides ample opportunities to understand evolutionary processes that shape biodiversity.

The genus *Arvicola* (Lacépède, 1799), commonly known as water voles, is an example of a genus of rodents that have puzzled researchers for many years. Water voles have a wide Palearctic distribution and occur as far west as the Atlantic coast, east to the Lena river, north as the Arctic coast, and south as the Northern Galilea in Israel as well as the Zagros mountains in Iran (Mahmoudi et al., 2020). Across their range, they display considerable morphological variation, and differences in the first lower molar (m1) have often been used for taxonomic classification of lineages (Desclaux et al., 2026; Escudé et al., 2008; Piras et al., 2012). Specifically, the SDQ (Schmelzband-Differenzierungs-Quotient) ratio, a measurement of the relative thickness of the enamel between the anterior (leading) edge, and the posterior (trailing) edge in the dental triangles of the m1, has been used to infer evolutionary relationships (Heinrich, 1978; Kalthoff et al., 2007; Koenigswald and van Kolfschoten, 1996).

Water voles, which have rootless (arhizodont) molars are believed to have evolved from *Mimomys savini* which had rooted (rhizodont) molars, during the early Middle Pleistocene, with the *Mimomys savini–Arvicola mosbachensis* transition, generally dated to approximately 700–600 ka in Central Europe (Gromov and Ya Polyakov, 1992; Maul et al., 2014; Maul and Markova, 2007). Water voles from the Iberian Peninsula and Iran generally have high SDQ-values similar to *M. savini* (“primitive”), while water voles from Northern and Central Europe have lower SDQ-values (“derived”) (Mahmoudi et al., 2020; Masini et al., 2020). This pattern is believed to have evolved through postglacial recolonisation from Iberian, Southwest Asian, and Northeastern continental refugia and subsequent diversification (Brace et al., 2016; Chevret et al., 2020; Mahmoudi et al., 2020; Piertney et al., 2005). Water voles from Italy have slightly lower SDQ-values than populations in the Iberian Peninsula and Iran, but still more ancestral enamel patterns than water voles from Northern and Central Europe (Mahmoudi et al., 2020; Masini et al., 2020), due to long-term isolation within the Italian Peninsula (Solano et al., 2024). In addition, there are two morphotypes of water voles, aquatic and fossorial (Kryštufek et al., 2015), which further contributes to the difficulties of taxonomic assignment based on morphology.

In recent years, genetic studies have improved our understanding of evolutionary relationships within *Arvicola*. Currently, four species are recognized in the Mammal diversity database (Burgin et al., 2018); *A. amphibius* (European, Siberian and Anatolian lineages), *A. sapidus* (Iberian Peninsula), *A. persicus* (Iran), and *A. destructor* (Italian Peninsula, Switzerland and Slovenia), hereafter referred to as *Arvicola italicus* following G.S Miller (Miller, 1910). However, discussions are still ongoing regarding their taxonomic classification. Chevret et al. (2020) proposed that there are three species within the genus; *A. amphibius*, *A. persicus*, and *A. sapidus*, Kryštufek and Shenbrot (2022) recognised four, additionally including *A. italicus*. Other authors have argued that the Swiss lineage, *A. monticola*, should also be considered as a distinct species (Mahmoudi et al., 2020; Pardiñas et al., 2017). Here, we follow this nomenclature and refer to samples from northern UK and one sample from Switzerland as belonging to the *A. monticola* lineage.

Mahmoudi et al. (2020) first discovered that water voles from Iran, although morphologically similar to *A. amphibius*, belong to a highly diverged lineage that split from the Eurosiberian lineage ca 2.5 million years ago (Ma). Subsequently, phylogenetic analyses of the mitochondrial cytochrome b (cytb) gene suggest that *A. sapidus* branched off ca 1.5 Ma, followed by the divergence of *A. italicus* from *A. amphibius* + *A. monticola* ca 861 thousand years ago (ka), and finally by the split between *A. amphibius* and *A. monticola* around 671 ka. In addition to this, major divergences evolved between the European and Siberian *A. amphibius* ca 484 ka and between northern and southern *A. italicus* ca 441 ka. However, analyses of the nuclear interphotoreceptor retinoid binding protein (irbp) gene revealed inconclusive results regarding interspecies relationship within the genus, for example failing to detect genetic structure between *A. amphibius* and *A. monticola* (Mahmoudi et al., 2020).

Previous attempts to genetically resolve evolutionary relationships within the genus *Arvicola* have been based on single mitochondrial and nuclear genes (Chevret et al., 2020; Kryštufek et al., 2015; Mahmoudi et al., 2020). However, phylogenies based on single genes often overestimate divergence times, especially in fast evolving taxa (Nichols, 2001). Furthermore, previous comparisons of mitochondrial and nuclear phylogenies indicate discrepancies between the two, but this could not be fully resolved using single genes (Mahmoudi et al., 2020). Therefore, in this study we set out to further investigate the evolutionary history of the genus *Arvicola* by analysing entire mitochondrial genomes, as well as whole genome data from specimens from all currently known lineages within the genus. Notably, our molecular dating reveals substantially more recent divergence times than previously reported. We also further characterise a recently identified lineage closely related to *A. persicus*, reported from Azerbaijan and here shown to occur also in Georgia and Anatolia, which we refer to as the *ognevi* lineage following the taxonomic treatment of Ognev (1950), and which we estimate to have diverged from *A. persicus* ca. 216 ka.

## 3 Materials and Methods

### 3.1 DNA extraction and library preparation

Ancient MI-labelled samples were processed in a clean room type laboratory at the University of Warsaw Laboratory of Paleogenetics and Conservation Genetics, Warsaw, Poland, while modern WM-labelled samples were processed in a semi-clean room type laboratory. The ancient AL-labelled samples were processed in the ancient DNA (aDNA) lab at the Centre for Palaeogenetics, Stockholm, Sweden, while the more recent AL-labelled museum specimens were processed in a part of the lab dedicated to historical specimens. Standard aDNA laboratory procedures were followed continuously to minimize the risk of contamination.

### 3.2 Modern samples

We examined a total of 89 modern samples. WM-labelled specimens consisted of 56 ethanol-preserved tissue samples, three molars, one bone sample (mandible) and six dried-up tissues (toes with nails) (Table S1). AL-labelled specimens were collected from the Museum of Natural History, Stockholm, Sweden, and from the Museum Koenig Bonn, Bonn, Germany, as well as from field work in Bulgaria (Tundzha) and Italy (Marina di Pisa). AL-labelled samples consisted of four molars and/or parts of mandibles, and 20 incisors.

For WM-labelled samples, dried-up tissues were rehydrated and cut in smaller pieces prior to extraction. Ethanol-preserved and dried-up tissue samples were extracted using Qiagen DNeasy Blood & Tissue Kit according to manufacturer’s instructions. Bone and teeth samples were pre-treated with 0,5% hypochlorite, crushed into smaller pieces and lysed overnight prior to DNA extraction, which was performed following Rohland et al. (2018). Double-stranded, double-indexed libraries were prepared as described in Meyer and Kircher (2010). Amplified libraries were purified, pooled and sequenced using paired-end 150 bp mode (PE150) on Illumina NovaSeq X Plus platform.

AL-labelled samples were crushed into smaller pieces using a pestle, and bleach-treated prior to DNA extraction as previously described. Lysis was performed at 55℃ for 24-48 hours, until samples were dissolved. DNA was extracted following the methods described in Dehasque et al. (2022), excluding pre-digestion. Double-stranded, double-indexed libraries were prepared from the extracts following Meyer and Kircher (2010) and treated with uracil-specific excision reagent enzyme (USER) as described in Dehasque et al. (2022). Initial low-depth sequencing was performed for all samples on an Illumina NovaSeq X Plus platform with a 2x150 bp setup at the National Genomics Infrastructure Sweden (NGI), Stockholm. Five samples (AL287, AL387, AL392, AL405 and AL413) were sequenced to higher coverage with the same setup as during initial low-depth sequencing at NGI, Stockholm. Further details regarding laboratory methods can be found in Supplementary Methods 2.1.

### 3.3 Ancient samples

Seven ancient samples, consisting of molars and/or mandibles, were included (Table S2). For MI-labelled samples, DNA was extracted following Dabney et al. (2013). Single-stranded, double-indexed libraries were prepared following the protocol of Gansauge et al. (2017). The AL-labelled samples were subjected to bleach treatment as described in Lord et al. (2022), modified from Boessenkool et al. (2017) prior to DNA extraction. Lysis was done as in Dabney and Meyer (2019), and DNA extraction was performed on a 96-column plate using 100 µl of lysate as described in Gilardet et al. (2025). Single-stranded, double-indexed libraries were built following Kapp et al. (2021), as modified by Nguyen et al. (2023), and cleanup was performed on a 96-column plate. Size selection was performed using Sera-Mag™ beads (Cytiva). Five samples, two from Łabajowa cave (MI3095 and MI3096), two from Trou Al’Wesse (AL265 and AL266) and one from Dzudzuana cave (MI737) were shotgun sequenced while two samples from Nad Tunelem cave (MI107 and MI109) were enriched for mitochondrial DNA as described in Baca et al. (2023). Libraries were sequenced on NextSeq550 and NovaSeq6000 in paired-end 50 bp mode (PE50), or on a NovaSeq X Plus platform with a 2x150 bp setup.

### 3.4 Processing of sequencing data

For all modern samples, except AL-labelled samples, AdapterRemoval v.2 (Schubert et al., 2016) was used to trim and merge the raw data. The filtered reads were then mapped to the mtDNA sequence of *A. amphibius* (MN122834, (Margaryan et al., 2021) using *bwa mem* with default parameters. Trimming and competitive mapping of AL-labelled samples was done using DNAharvester v.0.2.1 (Sharif et al., 2026). Filtered and merged reads were mapped to the *A. amphibius* reference genome mArvAmp1.2 (NCBI RefSeq: GCF_903992535.2) concatenated with a mitochondrial sequence from *A. amphibius* (accession number MN122828.1), and the human reference genome GRCh38.p14 (NCBI RefSeq: GCF_000001405.40). Further details regarding BAM-file processing are provided in Supplementary Methods 2.1. The mitogenome scaffold was extracted from each filtered BAM-file with *samtools view*, and consensus sequences generated with ANGSD v.0.940 using the following settings: most common base (doFasta 2), base quality filter 30 (minQ 30), mapping quality filter 25 (minmapQ 25), removing duplicates and secondary reads (remove_bads 1), and calling only positions with a depth of at least 3X (setMinDepth 3). Ancient samples labelled “MI” were mapped using bwa backtrack algorithm with parameters optimised for aDNA (−o 2 −n 0.01 −l 1024). The reference sequence was merged with the human mitochondrial reference sequence in order to filter presumptive contamination. Only reads longer than 30 bp and with MapQ >25 was retained. Presumed duplicates were removed using *samremovedup.py* script (https://github.com/pontussk/samremovedup). Consensus sequences were called using a custom script based on *bcftools* v.1.9 *mpileup* (Danecek et al., 2021) and *ivar* v.1.3 *call* (Grubaugh et al., 2019). We called only positions with coverage higher than 2 and if a base was supported by more than 75% of the reads. Visual verification of alignment files was performed in Tablet (Milne et al., 2013). DNA damage pattern and length distribution of DNA molecules was evaluated using MapDamage v.2.08. (Jónsson et al., 2013). The mitochondrial genomes of the two ancient samples labelled “AL” were assembled through the iterative assembly workflow in DNAharvester v.0.2.1 (Sharif et al., 2026). After trimming and merging of raw FASTQ-files, Mapping Iterative Assembler (MIA) (Green et al., 2008) was used to reconstruct mitogenomes, using the *A. amphibius* mtDNA sequence MN122828.1 as seed reference genome. The final consensus sequences were generated applying a minimum read depth of 3x, base quality of 40, and allele consensus agreement of 66%, using a custom python script implemented in DNAharvester (https://github.com/aersoares81/mia-helper-scripts).

### 3.5 Radiocarbon dating

Two samples (MI107 and MI109) were directly AMS radiocarbon dated at the Poznań Radiocarbon Laboratory, Poland, while radiocarbon dates for four additional ancient samples were obtained from published sources. All radiocarbon dates were calibrated in OxCal v. 4.4 (Ramsey and Lee, 2013) using the IntCal20 (Reimer et al., 2020) calibration curve.

### 3.6 Phylogenetic analyses of mtDNA

We added an additional 18 published mitochondrial genomes from modern samples to our dataset. These included one sample from Russia (MT381921), one sample from Israel (PP291566), two samples from Denmark (N122828 and MN122834), and 14 samples from the UK (UK584-UK597). In total, the dataset consisted of 115 mitochondrial sequences; 108 sequences from modern specimens, and seven sequences from ancient specimens. The sequences were aligned using MAFFT v. 7.526 (Katoh and Standley, 2013), and the final length of the mtDNA alignment was 16054 bp.

The best partitioning scheme and suitable substitution models were identified using PartitionFinder2 (Lanfear et al., 2017) (Table S3). Phylogenetic analyses were performed using BEAST v.1.10.4 (Suchard et al., 2018), with a strict clock applied to all partitions, and a Bayesian SkyGrid model, which were the most suitable settings according to Bayesian evaluation of the temporal signal (BETS) (Supplementary Text 2.2). Five radiocarbon dated samples were used to calibrate the molecular clock (Table S2). In addition, two samples with unknown ages were included, MI3096 and MI737. A uniform prior (initial value= 68,000, upper= 150,000, lower= 50,000) was applied to MI3096 which was estimated to have an infinite radiocarbon date. A uniform prior (upper= 50,000, lower= 0) was also applied to MI737. Three independent trees running for 100 M generations each were generated, and each run was inspected in Tracer v.1.7.2 for convergence. The trees were then combined with LogCombiner v.1.10.4 applying 10% burn- in and a resample frequency of 20,000. A consensus tree was generated from 13,501 trees using TreeAnnotator, and visualised using FigTree v1.4.4 (Rambaut, 2018). Further details are provided in Supplementary Text 2.2.

### 3.7 Mapping of nuclear DNA and reference filtering

Filtered reads were mapped to mArvAmp1.2 reference genome (GCF_903992535.2) using *bwa mem*, deduplicated using *samtools markdup* and realigned around indels using GATK 3.8 IndelRealigner (McKenna et al., 2010). Guided by mismatch analysis from AMBER (Dolenz et al., 2024) we discarded all reads shorter than 50 bp or 80 bp to minimize reference bias.

We used RepeatModeler2 (Flynn et al., 2020) to create a library of repetitive regions in the mArvAmp1.2 assembly and masked those fragments using RepeatMasker. We also ran RepeatMasker using freely available Dfam libraries from rodents (species −Rodentia). The combination of these two approaches resulted in 943 Mb of the reference genome masked. In addition, we used GenMap (Pockrandt et al., 2020) with -K 100 -E 2 arguments to assess mapability across the reference and excluded all sites with mapability score lower than 1. This analysis yielded 119 Mb. We retained only autosomal chromosomes; consequently, all genome-based analyses were conducted using 1,272 Mb, corresponding to 59.2% of the reference genome.

### 3.8 Variant calling

Ten samples generated in this study and additional 14 samples from the UK downloaded from ENA repository yielded mean genome coverage above 12 and for those samples we called whole genome VCFs using *bcftools* v.1.9 *mpileup* and *call* commands. We filtered out positions covered by less than 10 or more than 50 reads, positions called with quality below 20. We removed heterozygotes if one of the alleles were supported by less than 20% of the reads covering the position in question. Finally, we removed all the indels and SNPs within 5 bp from indels.

For the whole dataset of 45 samples the genotype likelihoods (GL) were called using ANGSD 0.930 (Korneliussen et al., 2014). We used the following settings: -minMapQ 25 - minQ 25 -uniqueOnly 1 -remove_bads 1 -minFreq 0.05 -minInd 34 -GL 2 -doGlf 2 - doMajorMinor 1 -doMaf 2 -SNP_pval 2e-6 -doCounts 1 -makeMatrix 1 -doCov 1. This resulted in a dataset of ca. 24 million variable positions. In addition, we computed an identity- by-state (IBS) distance matrix using -doIBS 1 argument in ANGSD on a dataset supplemented with the high coverage genome of *Ellobius lutescens* (ENA accession: PRJNA305123), serving as an outgroup.

### 3.9 Population structure and demographic history

We used the IBS distance matrix to reconstruct a neighbour-joining tree using the *ape* package in R (R Core Team, 2021). Node support was assessed using a block bootstrap approach. The covariance matrix based on GLs was visualised using Principal Component Analysis (PCA) computed with *eigen* command in R and plotted using ggplot2. See Supplementary Text 2.3 for more details.

The admixture proportions were inferred using NGSadmix (Skotte et al., 2013) based on GLs. To reduce linkage among sites and computational burden, we thinned the original Beagle file by retaining every 10th position, resulting in approximately the 2.3 million sites used in the analysis. We ran 30 replicates for each K from 2 to 12. We considered a given K to have converged when the difference in log-likelihood among the five highest-scoring replicates was <2. If this criterion was not met, an additional 30 replicates were run, and the replicate with the highest log-likelihood was retained. Runs for K= 2–9 met the convergence criterion. Model fit for the inferred admixture proportions was evaluated using evalAdmix v0.962 (Garcia-Erill and Albrechtsen, 2020).

To estimate the timing of population splits, we applied the Two-Two (TT) method to high-coverage genomes (Sjödin et al., 2021). First, we reconstructed ancestral states across the genome using data from three outgroup species, following the recommendations of the method’s authors. We generated a high-coverage genome of *Stenocranius gregalis* from the southern Urals (25× mean genomic coverage; accession: ERS30593375) and used the previously described genomic dataset for *E. lutescens*, together with publicly available genomic data for *Clethrionomys glareolus* (30× mean genomic coverage; accession: SRR4014963). All three outgroups were mapped to the mArvAmp1.2 reference genome. Whole-genome FASTA sequences were then generated for each species using ANGSD with the -doFasta 2 option. Ancestral-state files were constructed by assigning a nucleotide at sites where at least two of the three outgroups carried the same allele; all remaining sites were coded as N. Population split times were subsequently estimated following the workflow provided in the TT-method GitHub repository (https://github.com/jammc313/TT-method).

We performed pairwise sequential markovian coalescent (PSMC) analysis on medium and high coverage genomes (Li and Durbin, 2011). We generated PSMC input files using vcfutils.pl vcf2fq on filtered whole genome vcf files. We ran PSMC analyses using the atomic interval pattern originally proposed for human datasets (4+252+4+6). We generated 100 bootstrap replicates to evaluate the support for the inferred demographic trajectories. We scaled the TT divergence-time estimates and PSMC effective population size (Ne) trajectories assuming two generations per year and a per-site mutation rate of 5 × 10⁻⁹ substitutions per generation, estimated for the house mouse (*Mus musculus*; (Uchimura et al., 2015). Additionally, we repeated the scaling using a lower mutation rate of 2.96 × 10⁻⁹ substitutions per site per generation estimated for the brown rat (*Rattus norvegicus*) (Deinum et al., 2015).

To investigate the more recent history of the water voles from the UK Aberdeen population we first identified potentially related individuals using ngsRelate (Korneliussen and Moltke, 2015). This identified potential second-degree relatedness in four pairs of samples, so we removed three of those samples with the lowest coverage keeping seven for downstream analyses. We generated site allele frequency likelihoods (SAF) using ANGSD and converted it into folded and unfolded site frequency spectra (SFS) using realSFS (Korneliussen et al., 2014). The resulting SFS were used as an input for StairwayPlot2 (Liu and Fu, 2020) (See Supplementary Information Text 2.4–2.5 for more details).

### 3.10 Gene flow

To investigate the gene flow between water vole lineages and populations we used TreeMix v1.13 (Pickrell and Pritchard, 2012). For this analysis, individuals were assigned to population groups based on their placement in the mitochondrial and nuclear phylogenies. We used two alternative grouping schemes. In the first, individuals were divided into 11 groups representing the major mtDNA and nuclear lineages: *A. persicus*, *Georgian ognevi*, Anatolian *ognevi*, *A. sapidus*, *A. italicus*, *A. monticola*, four phylogeographically distinct groups within *A. amphibius* and the *Ellobius* as an outgroup. In the second scheme, *A. amphibius* individuals were combined into two groups, resulting in eight ingroup populations in total (Table S1).

Following the approach of Garcia-Erill et al. (2024) we estimated SAF separately for each group using ANGSD. SAF files from all groups were then intersected to retain only sites represented in every group. For each retained site and population, the allele count corresponding to the maximum-likelihood allele frequency was used to generate the input for TreeMix. This approach allows allele-count information to be derived directly from genotype likelihoods rather than from genotype calls, which is particularly suitable for datasets containing individuals with heterogeneous sequencing depth. We thinned the resulting TreeMix file tenfold using a custom script, resulting in approximately 5 million polymorphic sites. We ran 10 replicates of TreeMix analyses for each number of migration edges from 0 to 5 and selected the replicate with the highest likelihood for visualization. The optimal number of migration edges was inferred from the second-order rate of change in the likelihood (Δm), analogous to the Evanno method (Evanno et al., 2005), as implemented in OptM (Fitak, 2021). See Supplementary Text 2.6 for more details.

For the set with eight ingroup populations we reconstructed the admixture graph using AdmixtureBayes (Nielsen et al., 2023). We used the input file generated as in TreeMix analysis. We performed six independent runs for 30 million iterations and used 32 MCMC chains each (–n 600,000 and –MCMC_chains 32). We evaluated the convergence using the R script provided by the authors (https://github.com/avaughn271/AdmixtureBayes) and plotted three topologies with the highest posterior probabilities and best estimates of the branch lengths and admixture proportions.

We used *Dsuite* (Malinsky et al., 2021) on a dataset comprising 10 medium-high-coverage ingroup genomes, representing the main *Arvicola* lineages, and *Ellobius* outgroup. Individual vcf-files were merged and only biallelic sites were retained. The resulting multisample vcf was filtered and LD pruned using PLINK1.9 (Purcell et al., 2007), then f-statistics were calculated using *Dtrios* program and derived *f*-branch statistics were calculated using Fbranch command and the tree topology based on IBS NJ tree.

To assess whether post-divergence gene flow between Iranian *A. persicus* and Georgian *ognevi* could bias divergence-time estimates obtained with the Two-Two (TT) method, we performed demographic modelling in *fastsimcoal* v.2.8 (Excoffier et al., 2021). Analyses were based on the joint two-dimensional SFS constructed from three Georgian *ognevi* and four Iranian *A. persicus* genomes. We compared four alternative demographic scenarios: (i) strict isolation following population divergence, with no subsequent gene flow; (ii) early gene flow occurring during an initial period following divergence; (iii) a single pulse of admixture between the two populations; and (iv) continuous gene flow beginning at a specified time after divergence and continuing until the present. For each model, demographic parameters, including population divergence time, effective population sizes, and, where applicable, migration rates and the timing of gene flow, were estimated by composite-likelihood optimization. Multiple independent optimization runs were performed for each scenario, and models were compared using their maximum composite likelihoods and Akaike information criterion (AIC). See Supplementary Text 2.7 for more details.

### 3.11 Heterozygosity and Runs of Homozygosity (ROH)

We used ANGSD 0.930 to compute per sample heterozygosity for all samples with coverage above 5×. First, we calculated SAF (−dosaf 1), using the genotype likelihood algorithm from GATK (-gl 2) and next we used realSFS to generate a folded SFS in 20Mb windows. The heterozygosity was then inferred as the number of heterozygous sites divided by all sites.

Runs of homozygosity were inferred using two approaches. For medium to high coverage samples we used PLINK v1.9 –homozyg command (Purcell et al., 2007). The merged VCF was converted to PLINK format, retaining only biallelic SNPs and without applying a missing-data filter. Because the resulting dataset was extremely dense, with approximately one SNP every 50 bp, it was thinned to approximately one SNP every 2 kb, resulting in ∼900,000 SNPs. This was done to obtain a marker density more comparable to SNP-array datasets, for which the PLINK -*-homozyg* approach was originally developed. See Supplementary Text 2.8 for PLINK v1.9 arguments used for ROH discovery. Secondly, we used ROHan (Renaud et al., 2019) on all samples with mean coverage above 5× to infer heterozygosity and ROHs directly from read alignments. We used a window size of 500kb and a TSTV ratio of 2.21 estimated from high coverage genomes. See Supplementary Text 2.8 for more details.

To investigate potential consequences of low genetic diversity in water vole populations from the UK we assessed mutational load using SnpEff v.5.2 (Cingolani et al., 2012). The genome annotation of the water vole reference genome (mArvAmp1.2) was used to build a database for SnpEff. The rest of the workflow was performed as in (Lindahl et al., 2026). Variants were annotated into three different impact classes; low (non-disruptive, synonymous mutations), moderate (non-synonymous, but moderately disruptive), and high (highly disruptive, for example frame-shift and loss of function-mutations) impact. SnpSift v.5.0e was used to extract relevant fields from the SnpEff annotated files, and custom code was used to count the number of non-reference (derived) alleles in homozygous and heterozygous states. Mutational load of the different impact classes was calculated as follows; (counts of variants, homozygous variants counted as two)/(counts of derived alleles, homozygous alleles counted as two). The results were plotted using ggplot v.4.0.3 (Wickham, 2016). See Supplementary Text 2.9 for more details.

## 4 Results

### 4.1 Dataset

We generated mitochondrial genome sequences from 108 modern and historical water vole samples collected across Eurasia, including 18 sequences obtained from public repositories and representing all previously recognised major lineages (Fig. 1A; Table S1). To calibrate the mitochondrial phylogeny, we additionally recovered at least 70% of the mitogenome from seven palaeontological specimens (Table S2), six of which were directly radiocarbon dated (Table S3). We also generated low-to high-coverage nuclear genomes (0.98×–31.4×; median 12.5×) from a subset of 44 recent samples, including 14 from public repositories as well as from one palaeontological specimen (Table S1–2).

**Figure 1.**
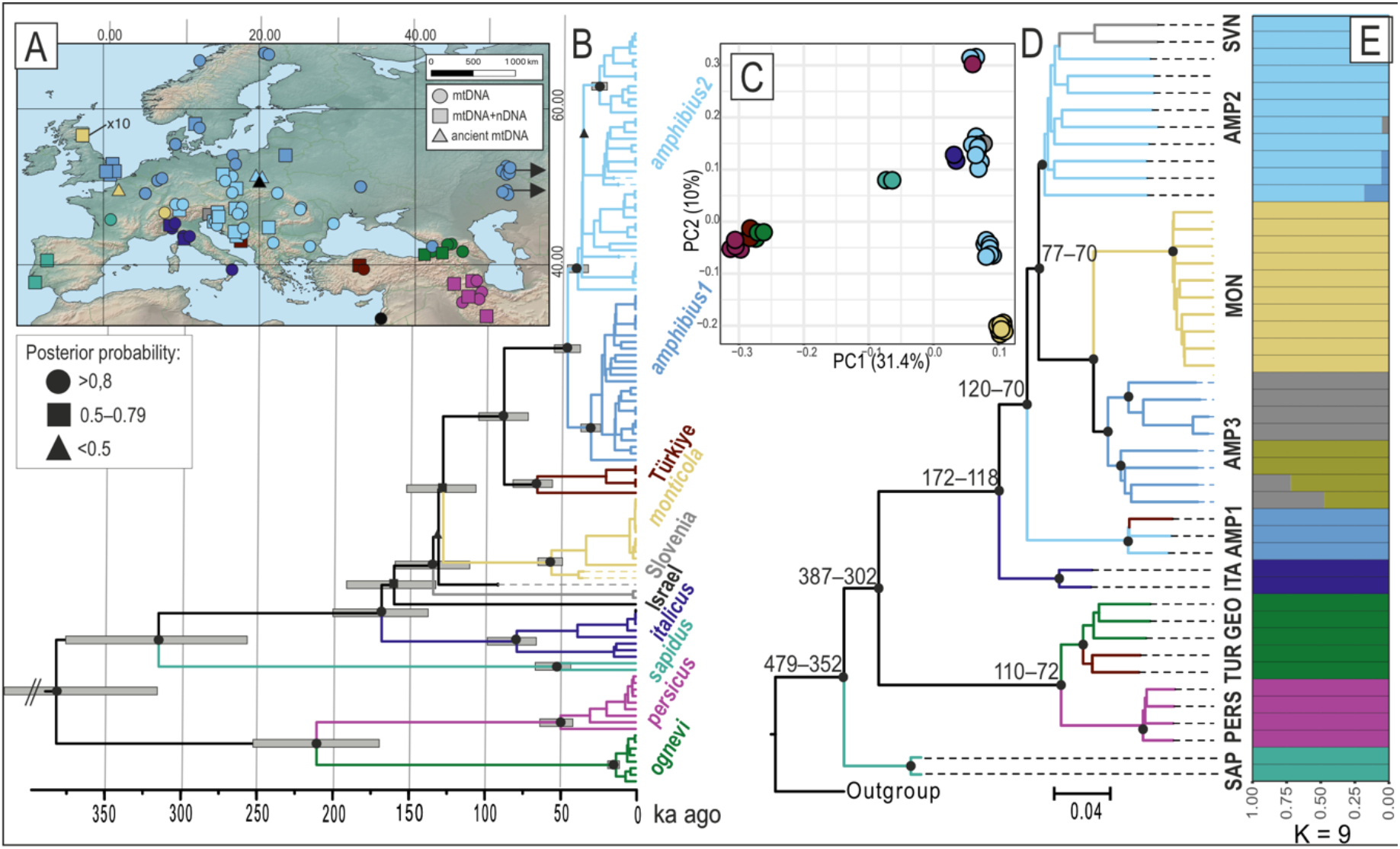
Phylogenetic relationships and population structure of Eurasian water voles. A) Sampling localities. Circles denote modern and historical samples for which mitochondrial DNA (mtDNA) sequences were obtained, squares denote modern and historical samples with both mtDNA and nuclear DNA data, and triangles denote ancient samples represented by mtDNA data. Samples are coloured according to the mitochondrial lineages shown in panel B. B) Time-calibrated mitogenomic phylogeny reconstructed in BEAST v1.10.4. Posterior probabilities are indicated by node symbols and are omitted for some younger nodes for clarity. Node bars represent the 95% highest posterior density (HPD) intervals for divergence times. The complete tree with tip labels is shown in Fig. S1. C) Principal component analysis based on genotype likelihoods, showing the first two principal components. Samples are coloured according to the mitochondrial lineages shown in panel B. D) Neighbour-joining tree of water voles based on identity-by-state distances calculated from nuclear genomic data. Branches are coloured as in the mtDNA phylogeny. Selected nodes are annotated with the minimum and maximum divergence-time estimates inferred using the Two-Two (TT) method. Dots at major nodes indicate 100% bootstrap support. The tree with tip labels is shown in Fig. S2. E) Admixture proportions inferred with NGSadmix assuming nine ancestral populations (K = 9). See Fig. S3 for admixture proportions inferred for K = 2–12.

### 4.2 Mitochondrial phylogeny

The topology of the reconstructed mitochondrial phylogeny (Fig. 1B) was generally consistent with previously published phylogenies based on mtDNA cytb sequences (Mahmoudi et al., 2020), but here we also obtained the lineage sister to *A. persicus* recently identified in samples from several localities in Georgia (Maglakelidze et al., 2024) and in three specimens from Azerbaijan (Bulycheva et al., 2025). We recovered several major mitochondrial clades and lineages: *A. persicus* (IRN) and its sister lineage, *ognevi*, from Georgia; *A. sapidus* (FRA and PRT); *A. italicus* (ITA); *A. monticola* (UK and CHE); a lineage represented by two specimens from Slovenia; another lineage represented by a single specimen from Israel; and *A. amphibius*, encompassing specimens from Western, Northern, Central and Eastern Europe, together with a distinct sister lineage represented by specimens from Türkiye. The time to the most recent common ancestor (TMRCA) of all sampled *Arvicola* lineages was estimated at approximately 383 ka (95% HPD: 315.8–459.9 ka), corresponding to the divergence of the lineage ancestral to *A. persicus* and the Georgian clade from the remaining Eurosiberian *Arvicola*. *A. persicus* and the Georgian lineage subsequently diverged from each other at approximately 216 ka (95% HPD: 169.6–252.6 ka). Within the Eurosiberian clade, *A. sapidus* was the first lineage to diverge, at approximately 315 ka (95% HPD: 256.7– 376.4 ka), followed by *A. italicus* at approximately 168 ka (95% HPD: 138.4–200.9 ka). *A. monticola* diverged from the lineage comprising *A. amphibius* and the Anatolian lineage at approximately 130 ka (95% HPD: 104.6–150.5 ka), whereas the split between the Anatolian lineage and *A. amphibius* was dated to approximately 89 ka (95% HPD: 70.6–108.3 ka). Several subsequent divergences were inferred within *A. amphibius*, including the split between a lineage currently distributed across Eastern, Northern and Western Europe (Amphibius 1) and a lineage occurring mainly in Central and Southern Europe (Amphibius 2).

### 4.3 Population structure and divergence

Analyses based on genome-wide nuclear data revealed a broadly similar pattern of genetic differentiation, but with several important differences from the mitochondrial phylogeny (Fig. 1C–E; Fig. S2–4). First, Portuguese *A. sapidus* formed the sister group to all other water voles in the nuclear NJ phylogeny. Second, specimens from Türkiye clustered with the Georgian samples in both the PCA and NJ phylogeny (Fig. 1C–D). Third, the UK *A. monticola* samples (MON) were nested within the broader nuclear diversity of *A. amphibius*, although they remained clearly differentiated from the other populations. Within *A. amphibius*, three main nuclear groups were apparent. The first comprised samples from Bosnia and Herzegovina (AMP1). The second included samples from Central Europe and the Balkans (AMP2); together, these two groups broadly corresponded to the Amphibius 2 mitochondrial group. The distinct Slovenian mitochondrial lineage was not differentiated in the nuclear analyses and clustered within the Central European/Balkan group. The third group included samples from Northern, Eastern and Western Europe, including southern UK specimens, and broadly corresponded to the Amphibius 1 mitochondrial group (AMP3).

Population split times estimated using the Two-Two (TT) method between pairs of high-coverage samples (Fig. 1D, Table S4) were broadly comparable in temporal scale to those inferred from the mitochondrial phylogeny. Although differences in topology precluded a direct comparison of all individual nodes, the two earliest splits in the nuclear phylogeny, namely the divergence of *A. sapidus* from all other *Arvicola* and the divergence of the *persicus– ognevi* lineage from the *amphibius–italicus–monticola* lineage, fell within similar time intervals to the corresponding early divergences inferred from the mitochondrial phylogeny, even though their order was reversed. Among the more recent divergences, the split of *A. italicus* and the diversification within *A. amphibius* closely matched the mitochondrial estimates, whereas the split between *A. persicus* and *ognevi* lineage (∼110–72 ka) was substantially younger than the corresponding estimate based on mtDNA.

### 4.4 Admixture and gene flow

We used several complementary approaches to investigate historical gene flow among water vole lineages and populations, including TreeMix analyses using two alternative population groupings, the *f*-branch statistic implemented in Dsuite and an admixture graph inferred with AdmixtureBayes (Fig. 2). These analyses produced partially overlapping signals of historical gene flow and admixture.

**Figure 2.**
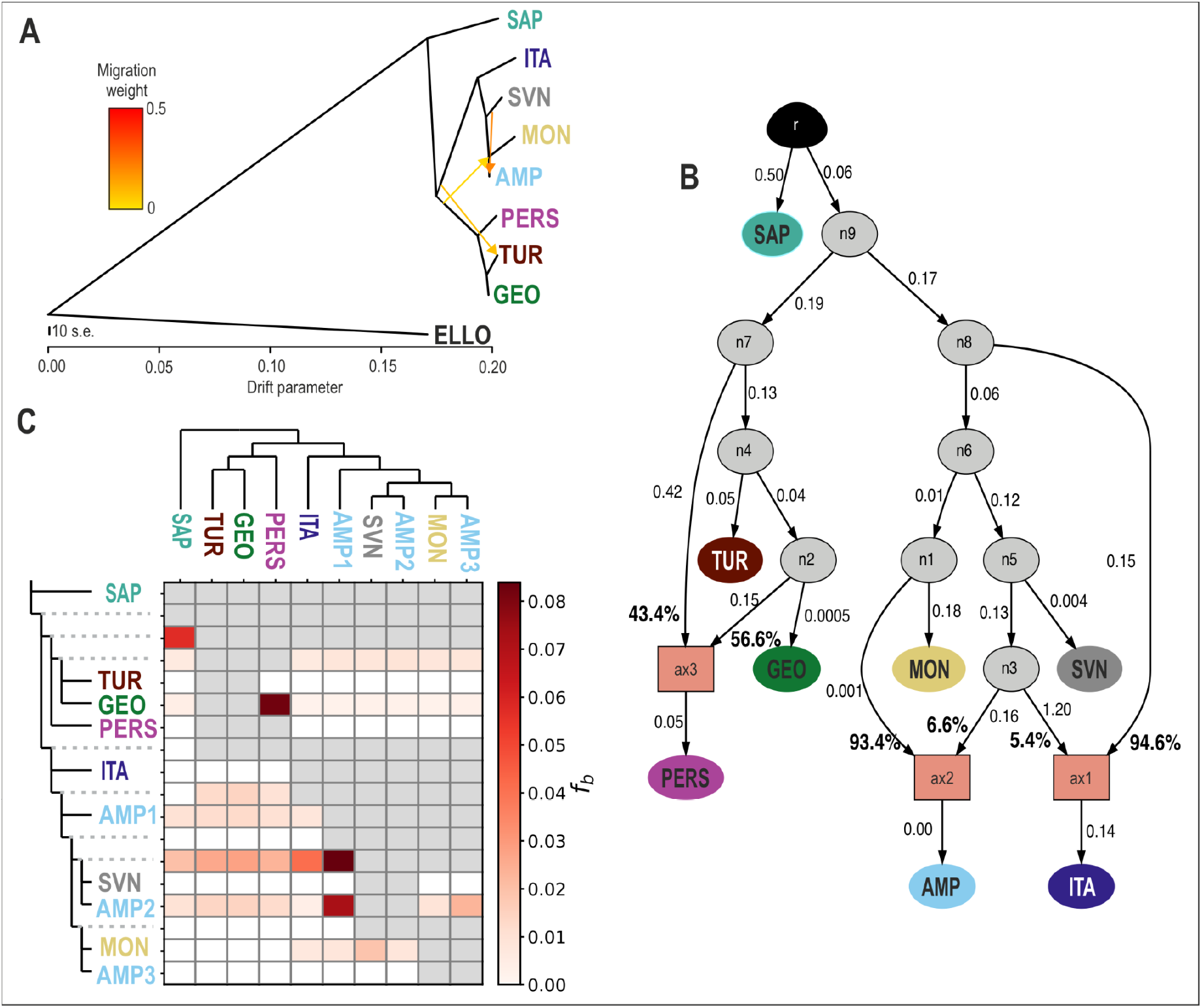
Patterns of gene flow and admixture among Eurasian water vole lineages. A) Maximum-likelihood admixture graph of water voles grouped into eight populations inferred with TreeMix with three migration edges allowed. Migration arrows are coloured according to their inferred weight. B) Admixture graph inferred with AdmixtureBayes, showing the relationships among the major lineages and inferred admixture events; percentages indicate admixture proportions. C) Pairwise *f*-branch statistics inferred with Dsuite summarizing signals of excess allele sharing among water vole lineages.

Both TreeMix analyses (Fig. 2A; Fig. S5, S7) inferred gene flow involving the lineage ancestral to *A. monticola* and northern *A. amphibius* (AMP3), with ancestry derived from a lineage ancestral to *A. italicus* and *A. amphibius*. This signal is consistent with the discordant position of *A. monticola* in the mitochondrial and nuclear phylogenies (Fig. 1; Fig. 2A).

A second signal involved the Anatolian samples (TUR), which showed pronounced mitonuclear discordance. In the mitochondrial phylogeny, TUR was sister to *A. amphibius*, whereas in all nuclear analyses, including the NJ phylogeny, PCA and ngsAdmix, the Anatolian samples clustered with Georgian *ognevi*. Consistent with a history of admixture between these lineages, the TreeMix analysis with eight ingroup populations inferred a migration edge from the lineage ancestral to *A. amphibius* and *A. italicus* towards TUR (Fig. 2A; Fig. S5–6).

Evidence of gene flow involving *A. italicus* was also recovered. In the admixture graph, *A. italicus* was modelled with a minor ancestry component derived from an ancestral lineage also contributing to *A. amphibius* (Fig. 2B). Consistently, the *f*-branch analysis showed excess allele sharing between *A. italicus* and the ancestral branch leading to the Balkan/Slovenian *A. amphibius* samples (Fig. 2C), while the more finely partitioned TreeMix analysis inferred a migration edge involving these lineages (Fig. S7–8).

The strongest and most consistently signal of gene flow involved *A. persicus* (PERS) and Georgian *ognevi* (GEO). These two lineages showed a high *f*-branch value (Fig. 2C), while the admixture graph modelled *A. persicus* as receiving nearly half of its ancestry from a lineage closely related to Georgian *ognevi* (Fig. 2B). The finer partitioned TreeMix analysis also recovered a reticulate relationship among the West Asian lineages (Fig. S7–8). The divergence genome-based time divergence-time estimate of *A. persicus* and Georgian ognevi was substantially younger than the mitochondrial estimate. To test whether extensive gene flow could have affected the nuclear divergence-time estimate of *A. persicus* and Georgian *ognevi*, we used *fastsimcoal2* to compare four demographic scenarios: strict isolation, ancient gene flow, a single pulse of gene flow and recent gene flow. Of these, the model with recent, continuous gene flow provided the best fit to the observed data, supporting continued genetic exchange between the two lineages after their initial divergence (Supplementary Text 2.7, Tables S6–7). This model inferred an ancient (>400 ka) divergence between *A. persicus* and Georgian *ognevi.* Thus, the comparatively young divergence inferred from the nuclear genomes by the TT method more likely reflects extensive gene flow after the split than a recent separation of the two lineages.

### 4.5 Demographic history, genetic diversity and load

We used the pairwise sequentially Markovian coalescent (PSMC) to reconstruct effective population size (N_e_) trajectories for medium- to high-coverage genomes (Fig. S9). Although demographic trajectories varied among lineages and populations, several general patterns were apparent. The southern lineages, including *A. sapidus*, *A. persicus* and *A. italicus*, generally showed comparatively low N_e_ throughout the Late Pleistocene, with relatively limited changes in *A. sapidus* and *A. persicus*. Georgian and Anatolian *ognevi* showed an increase during the Late Pleistocene followed by a decline towards the Holocene. Demographic trajectories within *A. amphibius* were more variable but all experienced an increase in the Middle Pleistocene. The Bosnian individual WM813 related to AMP1 cluster showed several fluctuations in N_e_ during the Late Pleistocene, whereas the Hungarian individual WM818 (AMP2) exhibited a comparatively steady increase towards the most recent part of the trajectory. The Slovenian individual WM798 from a distinct mtDNA lineage showed an increase to a maximum at approximately 40–30 ka, followed by a decline. Similar trajectories were observed in southern UK *A. amphibius*, although their maximum inferred N_e_ values were generally higher. Northern UK water voles belonging to the *A. monticola* mtDNA lineage showed an even more pronounced increase, reaching the highest N_e_ values among the analysed populations at approximately 50–30 ka, followed by a steep decline towards the Holocene.

To further investigate the recent demographic history of northern UK water voles, we applied StairwayPlot2 to seven unrelated individuals from the Aberdeen population. The analysis indicated an approximately fivefold decline in N_e_ around the Pleistocene–Holocene transition, consistent with the decline inferred from PSMC, followed by a second pronounced reduction around 4 ka and a subsequent recovery (Fig. S10). Both features were recovered using folded and unfolded site frequency spectra, showing that the demographic signal was not strongly dependent on ancestral-state polarization.

Genetic diversity of water voles was assessed using genome-wide heterozygosity and the burden of runs of homozygosity. Each measure was estimated using two complementary approaches, which yielded broadly consistent results (Fig. 3; Figs. S11–S12; Supplementary Text 2.8). Patterns of heterozygosity and ROH burden in modern samples partially reflected the Late Pleistocene N_e_ trajectories inferred with PSMC. Balkan *A. amphibius* (AMP1 and AMP2) showed the highest genome-wide heterozygosity and generally low ROH burden, a pattern also observed in Georgian *ognevi.* Anatolian *ognevi* showed slightly lower heterozygosity and a greater contribution of shorter ROHs (<16 Mb). The southern lineages, *A. sapidus*, *A. persicus* and *A. italicus*, generally showed lower heterozygosity, although only *A. sapidus* exhibited a marked increase in the contribution of longer ROHs, particularly in the 8–32 Mb size classes. The Russian *A. amphibius* samples (AMP3) were notable for particularly low heterozygosity, and one individual, WM825, had the largest fraction of its genome contained within long ROHs (>16 Mb). Southern UK *A. amphibius* (AMP3) also showed lower heterozygosity than Balkan *A. amphibius*, and one individual exhibited a substantial proportion of its genome in ROHs. In contrast, northern UK *A. monticola* consistently showed low heterozygosity together with a high ROH burden, with substantial fractions of the genome contained within ROHs.

**Figure 3.**
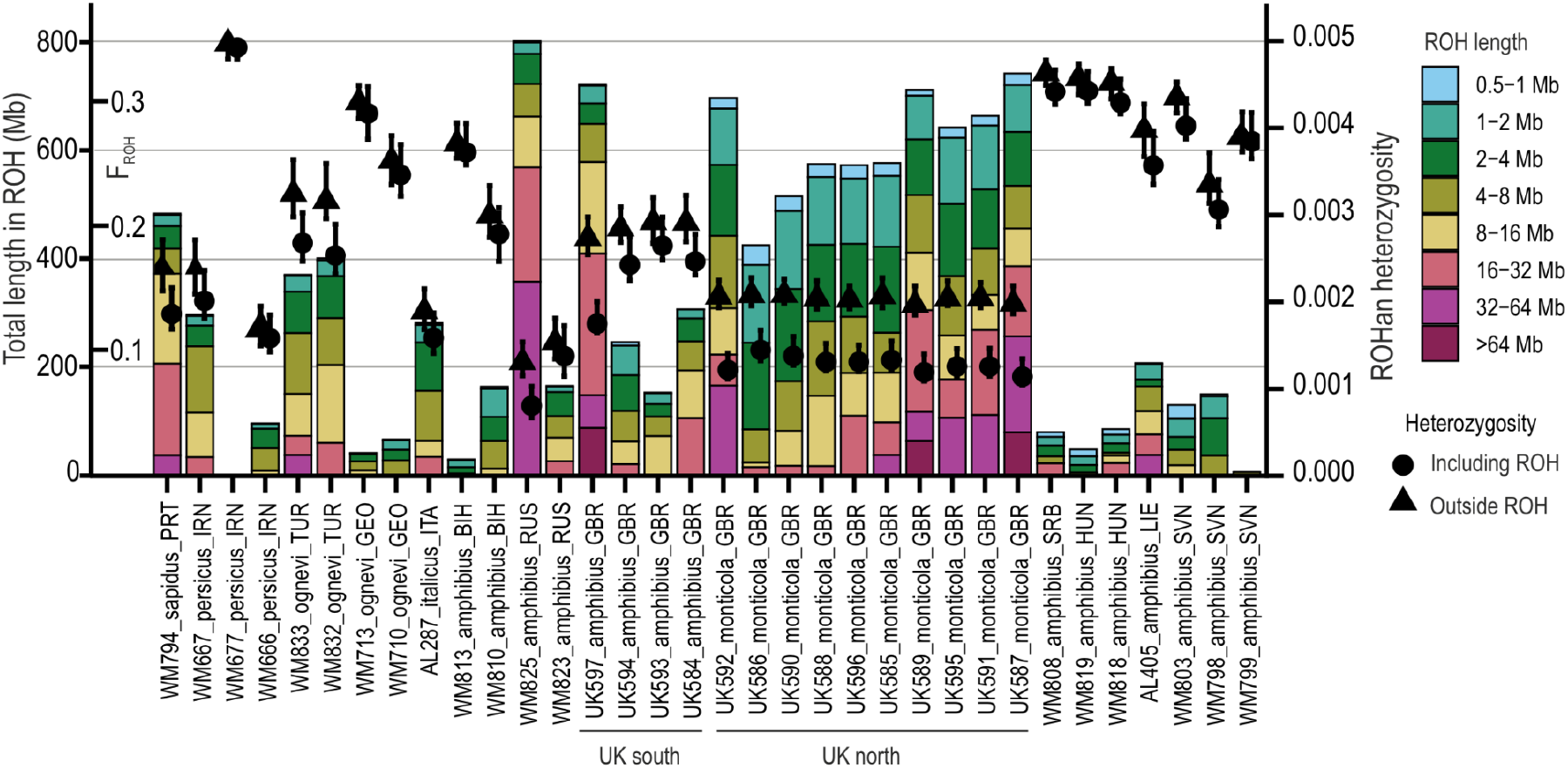
Runs of homozygosity and genome-wide heterozygosity inferred using ROHan for samples with mean genomic coverage >5×. Stacked bars represent the total length of ROHs, partitioned into ROH length classes. The corresponding F_ROH_ values were calculated as the summed ROH length divided by the total autosomal genome length (2.3Gb) and are indicated on the secondary scale of the left axis. Genome-wide heterozygosity (θ) is shown on the right y-axis; circles represent estimates including ROH regions and triangles estimates calculated outside ROHs. Error bars indicate the lower and upper ROHan estimates.

To assess potential consequences of low genetic diversity we estimated mutational load using SnpEff (Fig. S13). In addition to displaying high genetic diversity, Georgian *ognevi* also had low genetic load across all impact classes. Balkan *A. amphibius* had slightly higher genetic load in all impact classes in comparison, as did Anatolian *ognevi.* The southern lineages *A. sapidus* and *A. persicus* had similar genetic load in the high impact class, while *A. italicus* had the lowest genetic load in the high impact class of all samples (7.32 high impact mutations/100,000 derived alleles), but higher genetic load in the low impact class than *A. persicus* and *A. sapidus*. One sample *A. amphibius*, WM818 from Hungary, had relatively high genetic load across all impact classes. However, samples from southern UK *A. amphibius* (UK_SOUTH, AMP3), had the highest genetic load in the high impact class, and the sample with the most high impact mutations was UK593 with 12.58 high impact mutations/100,000 derived alleles (Table S10). Northern UK *A. monticola* (UK_NORTH), which displayed lower genetic diversity and higher ROH burden than southern UK *A. amphibius,* had lower genetic load in the high impact class compared to UK_SOUTH, but higher genetic load in the low impact class.

## 5 Discussion

The evolutionary history and taxonomic status of water voles has long been debated. By analyzing complete mitochondrial and nuclear genomes from samples representing all currently proposed water vole lineages, our study adds valuable new insights into the population structure and evolutionary history of the genus. While mitochondrial structure largely agrees with previous findings based on short genetic markers, with the addition of the Georgian lineage *ognevi* found to be closely related to *A. persicus,* nuclear DNA reveals previously unrecognized discordances. Furthermore, molecular dating suggests significantly younger divergence times than previously estimated, shifting the initial diversification of the genus to the Middle and Late Pleistocene rather than the Early Pleistocene.

The discrepancy between our divergence estimates and those of Mahmoudi et al. (2020) is quite large. For example, we estimate that *A. persicus* diverged from the remaining lineages ca 382 ka rather than 2.5 Ma, more than two million years later (Fig. 1B, Fig. S1). An important difference between the two analyses lies in the calibration strategy. Mahmoudi et al. (2020) estimated divergence times from mitochondrial cytb sequences using secondary calibration points of approximately 3 Ma for divergences among *Arvicola*, *Chionomys* and *Microtus*, whereas our mitochondrial timescale was calibrated directly using radiocarbon-dated Late Pleistocene *Arvicola* specimens. Such differences in calibration timescale may have a major effect on inferred divergence times because mitochondrial evolutionary rates are known to be time-dependent, with rates estimated over recent timescales often exceeding those inferred from deeper phylogenetic calibrations (Ho et al., 2015; Molak and Ho, 2015). The fossil record of water voles provides an independent perspective on these alternative timescales. In many arvicoline rodents, closely related evolutionary lineages show little or no morphological differentiation despite substantial genomic divergence. For example, the field vole species complex, comprising *Microtus agrestis*, *M. lavernedii*, and *M. rozianus*, includes genetically distinct lineages that are nearly impossible to distinguish in the fossil record (Baca et al., 2026; Navarro et al., 2018). In contrast, water voles exhibit well-documented temporal and geographic differentiation in dental morphology which allows molecular divergence estimates to be compared against the appearance of lineage-specific characters in the fossil record. For European populations, Koenigswald and Van Kolfschoten (1996) proposed SDQ thresholds distinguishing successive chronospecies *A. mosbachensis* (SDQ > 120), *A. mosbachensis/amphibius* (95 < SDQ < 120), and *A. amphibius* (SDQ < 95). Subsequent studies broadly supported a temporal decline in SDQ, with values generally falling below 120 between the Holsteinian and Saalian (ca. 380–300 ka) and below 100 from the Eemian onwards, reflecting an evolutionary trend from more primitive to modern morphologies (Heinrich, 1978; Kalthoff et al., 2007; Koenigswald and van Kolfschoten, 1996; Maul et al., 2014; Socha, 2014). However, Masini et al. (2020) emphasised that this scheme applies primarily within individual evolutionary lineages and that the history of *Arvicola* was more complex, particularly in the Iberian and Italian Peninsulas.

In this context, the Early Pleistocene *A. jacobaeus* from Atapuerca (∼1.2 Ma), proposed by Mahmoudi et al. (2020) as a possible ancestor of *A. sapidus*, was not recovered as a close relative of *A. sapidus* in its original description (Cuenca-Bescós et al., 2010). Also Maul et al. (2014) suggested that it may be more closely related to *Allophaiomys* than to the lineage leading to extant *A. sapidus.* Fossils showing clearer affinities to *A. sapidus* appear only during the Middle Pleistocene (Cuenca-Bescós et al., 2010).

Similarly, Italian *Arvicola* from Visogliano at ca. 400 ka still retain the primitive *A. mosbachensis*-like dental morphology, with differentiation of the Italian lineage becoming apparent considerably later, probably before the Eemian Interstadial (Masini et al., 2020; Maul et al., 1998). More generally, Maul et al. (2021) noted that regional morphological differentiation was limited in the early Middle Pleistocene but became much clearer from the late Middle Pleistocene onwards.

Although, as also noted by Mahmoudi et al. (2020), the morphological differentiation may postdate genetic divergence, the relatively late appearance of lineage-specific dental characters is, in our opinion, more consistent with the younger Middle to Late Pleistocene diversification inferred here than with the substantially older divergence times proposed previously.

Apart from the divergence times, we observe clear mitochondrial structure in our dataset, consistent with previous findings (e.g (Chevret et al., 2020; Mahmoudi et al., 2020). The earliest divergence, which separated *A. persicus* and the Georgian lineage *ognevi* from the remaining lineages, dates to MIS 12–MIS 11 (478–374 ka ago). During MIS 12–MIS11 there was a transition from an extensive glacial period to a long and warm interglacial (Candy et al., 2014; Shackleton, 1987), and the shifting climate may have affected terrestrial connectivity between Anatolia and Europe (Bulut et al., 2026), which could have contributed to the isolation and independent evolution of Caucasian water voles. The subsequent divergence of *A. sapidus*, which occurred during MIS 11–MIS 9 (424–300 ka ago), may have occurred through isolation on the Iberian Peninsula, which acted as an important glacial refugium for many species during climatic oscillations in the Pleistocene (Hewitt, 2004; Taberlet et al., 1998). Divergence of the remaining major lineages occurred throughout the late Middle and Late Pleistocene, beginning with *A. italicus* around MIS 7–MIS 6 (243–130 ka ago), followed by *A. monticola* during MIS (191–130 ka ago), although the support is relatively weak for this branch (posterior probability (PP) <0.95). Anatolian/Balkan water voles diverged from remaining *A. amphibius* around MIS 5 (∼130–71 ka ago), and finally we find that *A. amphibius* diverged into two subgroups during MIS 3 (∼57–29 ka ago), corresponding to lineages L1 and L2 in Chevret et al. (2020). It is clear that throughout the Pleistocene, water voles rapidly diverged into several distinct mitochondrial lineages, and several, though not all, of the divergence estimates coincide with those inferred for other rodents (Baca et al., 2023; Lord et al., 2022). Many voles and lemmings diverged during interglacials such as MIS 7 and MIS 5 likely due to expanding forests and fragmented landscapes. However, it is slightly more complex to pinpoint specific climatic events that may have driven divergence of water voles, possibly linked to the remarkable morphological and ecological plasticity of the genus, which may have triggered different responses to climate change compared to many other rodents (García-Rodríguez et al., 2024).

Although we find a similar mitochondrial structure to that reported in previous studies, nuclear data reveal several discordances with the mitochondrial data. In the PCA and NJ phylogeny, although most samples cluster according to mitochondrial haplogroup affiliation, there are several notable exceptions. Anatolian and Georgian samples cluster together and close to *A. persicus* (Fig. 1C) despite the placement of the Anatolian samples as sister lineage to *A. amphibius* in the mtDNA phylogeny. Furthermore, samples from Slovenia, which fall outside *A. amphibius* in the mitochondrial phylogeny, cluster with *A. amphibius* in the nuclear analyses. *A. monticola* also forms a distinct mitochondrial lineage but is nested within the broader nuclear diversity of *A. amphibius*. Together, these patterns point to a substantial role of historical gene flow in shaping the evolutionary history of *Arvicola*.

The Anatolian water voles have previously been treated as a subspecies of *A. amphibius* due to their placement in the mitochondrial phylogeny (Mahmoudi et al., 2020). In contrast, in our analysis of admixture proportions, samples from Georgia and Türkiye are modelled as a single component, even at K=9 (Fig. 1E). Furthermore, they appear to have a similar demographic history, with a decrease in N_e_ ∼120 ka ago, followed by an increase around ∼60– 70 ka ago, and a subsequent decrease into the Holocene (Fig. S9). This suggests that water voles from Türkiye actually belong to the *ognevi* lineage found in Georgia rather than *A. amphibius* but have acquired mitochondrial genomes through introgression with *A. amphibius* some time in the past. This is supported by the TreeMix population graph which shows gene flow from a lineage ancestral to *A. amphibius, A monticola* and *A. italicus (*AMP/ITA//SVN/MON) into Anatolian *Arvicola* (Fig. 2A). However, the signal is fairly weak, and for example not clearly picked up by admixture graph modelling (Fig. 2B), suggesting that gene flow occurred long ago or was limited. The mitochondrial divergence between Anatolian *Arvicola* and *A. amphibius* is estimated here to have occurred around the MIS 5/MIS 4 transition (∼89 ka ago) (Fig. 1B, Fig. S1). Around that time, the land bridge between Anatolia and Southeast Europe was re-established (Çağatay et al., 2016), which might have facilitated gene flow from *A. amphibius* in the Balkans into Anatolian *Arvicola*. Indeed, one sample from Bosnia and Herzegovina clusters closely with the Anatolian samples in the mitogenome phylogeny, supporting the theory of shared alleles between *Arvicola* populations in Southeast Europe and Türkiye (Fig. 1).

Attribution of the Anatolian samples to the *ognevi* lineage also provides a better fit with the morphological evidence, as Anatolian water voles show SDQ values that are considerably closer to those of *A. persicus* than to modern A. amphibius (Maul et al., 2021). A similar situation of mito-nuclear discordance may apply to the now largely extirpated Israeli water vole population (Marom et al., 2024). The only available historical specimen from Lake Hula carries a mitochondrial haplotype nested within European *Arvicola* diversity (Fig. 1), whereas morphologically this population shows closer affinities to *A. persicus* (Maul et al., 2021)

In addition to gene flow between Balkan *A. amphibius* and Anatolian *ognevi*, we detected excess allele sharing between *A. persicus* and the Georgian *ognevi*, between the two main *A. amphibius* groups, AMP1 and AMP2, and between AMP1 or *A. italicus* and an unsampled lineage ancestral to SVN and AMP2. Ultimately, this may partly explain the long-standing difficulty in defining clear taxonomic boundaries within the genus *Arvicola*.

In contrast with this evidence of gene flow and consistent with previous studies (Chevret et al., 2020; Mahmoudi et al., 2020), we find that *A. sapidus* is clearly distinct, and there is no support for admixture with other lineages in the recent past. This is possibly facilitated by pre- or postzygotic barriers caused by differences in chromosome numbers, as *A. sapidus* has n=40 chromosomes, while most other water voles have n=36 chromosomes (De La Guardia and Pretel, 1979). If this evolved early during speciation, it could have prevented admixture between *A. sapidus* and remaining water vole lineages early on, causing *A. sapidus* to evolve independently throughout much of the Pleistocene and Holocene. However, the taxonomic status of *A. monticola* and *A. italicus* can still be debated. Although *A. monticola* forms a monophyletic clade in the mitochondrial phylogeny, it is nested within *A. amphibius* in the nuclear phylogeny. This pattern also appears in the admixture graph modelling, which shows that *A. monticola* shares recent ancestry with *A. amphibius*. The only other available *A. monticola* specimen, the Swiss sample AL402, yielded very low nuclear coverage (0.04×), but nevertheless showed a closer relationship in the nuclear data to the geographically proximate

*A. amphibius* sample from Liechtenstein than to the UK samples (Fig. S14; Supplementary Text 2.3). Together, this suggests that gene flow between *A. monticola* and *A. amphibius* lineages may have occurred repeatedly across their range. Similarly, *A. italicus* appears diverged from *A. amphibius* in both the mitochondrial and nuclear phylogeny but shares excess alleles with a lineage ancestral to SVN and AMP2 and was modelled to have a 5.4% contribution from the *A. amphibius* lineage. In fact, *A. amphibius* has been found in Northern Italy, and it is possible that a hybrid zone exists, or has existed in the past (Solano et al., 2024). However, if a hybrid zone exists, gene flow appears to be limited. Chevret et al. (2020) calculated the genetic divergence between *A. monticola* and *A. italicus* and the other lineages and deemed them to not be divergent enough to be considered separate species from *A. amphibius*, although *A. italicus* may be undergoing a speciation process. We share this view, and in addition we propose that the *ognevi* lineage should be considered a subspecies of *A. persicus*. This was also proposed by Bulycheva et al. (2025), who classified water voles from Azerbaijan, which likely belongs to the *ognevi* lineage, as a subspecies of *A. persicus* based on morphology.

To further investigate the demographic and evolutionary history of Eurasian water voles, we estimated inbreeding levels as runs of homozygosity (ROH) and genome-wide heterozygosity, with a particular focus on water voles from the UK which are of conservation concern (McGuire and Morse, 2020). We find that inbreeding levels vary widely between different populations. *A. amphibius* populations from central Europe and the Balkans have low inbreeding and high genetic diversity, likely reflecting the gene flow occurring between populations of *A. amphibius*, as well as recent population expansions as seen for example in the demographic history of WM818 from Hungary, and WM813 from Bosnia and Herzegovina (Fig S9). More isolated and southern populations, such as *A. sapidus* and *A. italicus* have more short and intermediate ROHs (2–8Mb) ROHs and lower genetic diversity, and their demographic trajectories reveal a long history of low Ne. *A. persicus* also appears to have a history of long-term low Ne, but overall high genetic diversity and few ROHs. These differences may partly reflect the more restricted and fragmented distribution of suitable water vole habitat in southern parts of the range. In Anatolia and the Near East, water voles have disjunct and fragmented distribution, closely associated with permanent water bodies, streams, irrigation channels and marsh vegetation, and occur at relatively low densities (Kryštufek and Vohralík, 2005; Maul et al., 2021). *A. sapidus* is a habitat specialist which occupies patchy and scattered riparian and wetland habitats (Centeno-Cuadros et al., 2011). Italian water voles also exhibit patchy and fragmented distribution, and their populations are believed to decline (Castiglia et al., 2016; Solano et al., 2024). Thus, the low long-term N_e_ of the southern lineages may reflect lower continuity and availability of suitable habitat, resulting in smaller and more fragmented populations than in much of Central and Northern Europe.

However, southern populations of Eurasian water voles have higher genetic diversity than *Arvicola* in the UK, especially those belonging to the *A. monticola* lineage in northern UK. Previous studies have suggested that water voles colonized the UK in two separate waves around the Pleistocene–Holocene transition. First population carrying *A. monticola* mtDNA haplotypes appeared in England prior to the Younger Dryas (12.9–11.7 ka BP) likely during the Bølling–Allerød (∼14.7–12.9 ka BP) warm phase. resulting in the characteristic biogeographic pattern known as the “Celtic fringe” (Brace et al., 2016; Piertney et al., 2005; Searle et al., 2009).

Our genomic dataset suggests that these two colonisation waves were not completely independent. Northern UK water voles carrying *A. monticola* mtDNA haplotypes are nested within the *A. amphibius* diversity (Fig. 1D) and TreeMix analyses suggest the ancient admixture from the unsampled population above ITA/AMP/MON to the population ancestral to the northern *A. amphibius* and northern UK water voles. Such admixture could have occurred during secondary contact in Britain or earlier during the Late Pleistocene. The distinctive PSMC trajectories of the British water voles, exhibiting a pronounced increase during the Late Pleistocene, may also be consistent with this scenario as it was shown that population structure and admixture between differentiated population can produce such a pattern (Chikhi et al., 2018; Wang et al., 2020).

We observe a drastic decline in N_e_ in northern UK samples during the Holocene (Fig. S9). This pattern may reflect founder effects caused by the colonization of a few founders in the Holocene, but may also have been influenced by human induced landscape changes as agricultural practices in the UK intensified, which coincides with the bottleneck observed ∼3– 4 ka ago (Fig. S10) (Edwards et al., 2019, 2000; Woodbridge et al., 2014), and hydrological variability throughout the Holocene (Charman et al., 2006). In recent years, water voles in the UK have also been threatened by increasing habitat loss and fragmentation, and the introduction of the American mink (McGuire and Morse, 2020). We find that water voles in the northern UK have long ROHs and low genetic diversity compared to most samples from southern UK and other Eurasian water voles (Fig. 3), which may indicate that their populations are more fragmented and vulnerable. To investigate potential consequences of inbreeding we also estimated mutational load for mutations believed to either have a high, moderate, or low impact on the function of proteins (Fig S13.). Interestingly, samples from northern UK have fewer high impact mutations than samples from southern UK, a pattern previously attributed to purging of highly deleterious recessive alleles in populations exhibiting low diversity and high inbreeding such as Alpine ibex, mountain gorillas and kākāpō (Dussex et al., 2021; Grossen et al., 2020; Xue et al., 2015)

A useful comparison is provided by British bank voles (*Clethrionomys glareolus*), which show a similar Celtic-fringe phylogeographic pattern, with a genetically distinct northern population also sampled near Aberdeen (Kotlík et al., 2018; Searle et al., 2009). Despite this pronounced population structure, genome-wide heterozygosity was reported to be broadly similar among British bank vole populations (Kotlík et al., 2018). Thus, the particularly low diversity and extensive ROH observed in northern British water voles are unlikely to be explained by their Celtic-fringe ancestry alone and probably reflect additional, species- or population-specific demographic processes, potentially including more recent population decline and fragmentation.

However, although informative for conservation efforts of water voles in the UK, these estimates are only based on a single population of water voles from northern UK and should be considered with care. Future studies with wider geographic sampling and including ancient genomes as reference would be useful to further entangle the genetic consequences of the population declines in British water voles throughout the Holocene.

## 6 Conclusions

Our genomic analyses reveal a complex evolutionary history of *Arvicola*, reflecting Pleistocene diversification and subsequent gene flow. Molecular dating based on directly radiocarbon-dated ancient sequences places the initial diversification of modern water voles in the Middle Pleistocene, substantially more recently than previously proposed. Importantly, this younger timescale is concordant with both nuclear genome-based estimates and the fossil record. This agreement provides independent support for the use of radiocarbon-dated ancient sequences to calibrate molecular divergence estimates at this temporal depth and has broader relevance for studies estimating Pleistocene divergence times using ancient DNA tip calibrations. Genome-wide data further reveal several cases of mitonuclear discordance and recurrent admixture among geographically proximate populations, including a distinct *ognevi* lineage closely related to *A. persicus*, which also includes the Anatolian water voles. Demographic analyses also reveal pronounced variation in long-term effective population size and genetic diversity across the range, with several southern lineages showing lower N_e_ and genetic diversity, potentially reflecting more restricted and fragmented habitat availability, while northern UK population exhibit particularly low diversity and extensive homozygosity. Together, these results highlight the importance of integrating ancient DNA, genome-wide data and the fossil record when reconstructing evolutionary histories and evaluating taxonomic boundaries.

## 7 CRediT authorship contribution statement

**Aleksandra Deroń:** Conceptualization, Formal analysis, Investigation, Writing – original draft, Data curation. **Amanda Lindahl:** Conceptualization, Formal analysis, Investigation, Visualization, Writing – original draft, Data curation, Methodology. **Barbara Bujalska:** Project administration, Investigation, Writing – review & editing, Data curation. **Ahmad Mahmoudi:** Resources, Writing – review & editing. **Boris Kryštufek:** Resources, Writing – review & editing. **Armando Nappi:** Resources, Writing – review & editing. **Aleksander Bukhnikashvili:** Resources, Writing – review & editing. **Soraia Barbosa:** Resources, Writing – review & editing. **Damien Flas:** Resources. **Monika Knul:** Resources. **Boyan Milchev:** Resources, Writing – review & editing. **Karl Frafjord:** Resources, Writing – review & editing. **Miriam Belmaker:** Resources, Writing – review & editing. **Jan Decher:** Resources, Writing – review & editing. **Daniela C. Kalthoff:** Resources, Writing – review & editing. **John Stewart:** Resources, Writing – review & editing. **Jan M. Wójcik:** Resources, Writing – review & editing. **Emmanuel Desclaux:** Resources, Writing – review & editing. **Claudio Berto:** Resources, Writing – review & editing. **Adam Nadachowski:** Resources, Writing – review & editing. **Love Dalén:** Conceptualization, Funding acquisition, Supervision, Writing – review & editing. **Mateusz Baca:** Conceptualization, Formal analysis, Funding acquisition, Investigation, Supervision, Visualization, Writing – original draft, Data curation, Methodology.

## 9 Acknowledgements

This research was financed by the Polish National Science Centre grant 2020/39/B/NZ8/02853 to MB and the European Research Council Advanced Grant awarded to the project “PrimiGenomes” (ID: 101054984). We acknowledge computational resources provided by the National Academic Infrastructure for Supercomputing in Sweden (NAISS), funded by the Swedish Research Council. We are grateful to Giorgio Bardelli and Michela Podestà from the Museo Civico di Storia Naturale di Milano for granting access to some of the Italian samples.

## 10 Appendix X. Supplementary data

Supplementary data to this article consists of two files:

Supplementary Data 1: Contains Supplementary Methods and Results, Figures S1–S14 and Tables S5–S10

Supplementary Data 2: Contains Tables S1–S4

## 11 Data availability

Raw reads from shotgun sequencing and mapped reads from mtDNA target enrichment were deposited in the European Nucleotide Archive under the project accession number PRJEB130321. The alignment of the mtDNA sequences used in the phylogenetic reconstruction was deposited in the RepOD repository ( https://doi.org/10.18150/KZC2KW).

## 12 Declaration of generative AI and AI-assisted technologies in the manuscript preparation process

Generative AI tools were used to assist with script development, debugging, testing, and manuscript preparation. All AI-generated suggestions and text were critically reviewed and verified by the authors, who retain full responsibility for the content of the manuscript.

