## Supplementary_Data_1 for "Genomic insights into the evolutionary history and diversification of water voles (*Arvicola* spp.)"

Spis treści

### List of Supplementary Figures:

**Figure S1.** Tip-calibrated mitochondrial phylogeny of 115 samples of water voles (Arvicola sp.) made with BEAST v1.10.4

**Figure S2.** Neighbour-Joining tree of water voles based on identity-by-state distances calculated from nuclear genomic data

**Figure S3**. Admixture proportions inferred with NGSadmix for K = 2–12 ancestral components

**Figure S4**. Evaluation of the estimated admixture proportions using evalAdmix

**Figure S5.** Maximum-likelihood population graphs inferred using TreeMix for water voles divided into eight population groups

**Figure S6.** Selection of the number of migration edges in the TreeMix analysis for the dataset partitioned into eight population groups using OptM

**Figure S7.** Maximum-likelihood population graphs inferred using TreeMix for water voles divided into ten population groups

**Figure S8.** Selection of the number of migration edges in the TreeMix analysis for the dataset partitioned into eleven population groups using OptM

**Figure S9.** Demographic reconstruction of water voles using the pairwise sequentially Markovian coalescent (PSMC) model

**Figure S10.** Demographic history of the Aberdeen water vole population inferred using StairwayPlot2

**Figure S11.** Runs of homozygosity inferred using PLINK v.1.9 for samples with coverage >12×

**Figure S12.** Genome-wide heterozygosity estimated using ANGSD for samples with mean genomic coverage >5×

**Figure S13**. Mutational load in modern water voles (Arvicola sp.)

### List of Supplementary Tables

**Table S1.** Modern samples used in this study. MtDNA and nuclear mapping statistics – separate xlsx sheet

**Table S2.** Ancient samples used in this study. MtDNA mapping stats, and AMS radiocarbon dating results – separate xlsx sheet

**Table S3.** PartitionFinder2 best fitting partitioning scheme and substitution models for mitochondrial DNA – separate xlsx sheet

**Table S4.** Results of population split time estimation using TT method – separate xlsx sheet

**Table S5.** Results of Bayesian evaluation of the temporal signal and tree prior

**Table S6.** Parameters and search ranges of four demographic models analysed with *fastsimcoal2*

**Table S7.** Comparison of the four *fastsimcoal2* demographic models

**Table S8.** Total length of identified ROH segments (Mb) depending on the maximum number of missing genotypes per window.

**Table S9.** Effect of the *rohmu* parameter on the number and total length of runs of homozygosity (ROH) inferred with ROHan

**Table S10.** Number of high, moderate, and low impact mutations per 100,000 derived alleles inferred from SnpEff.

### 1 Supplementary Figures

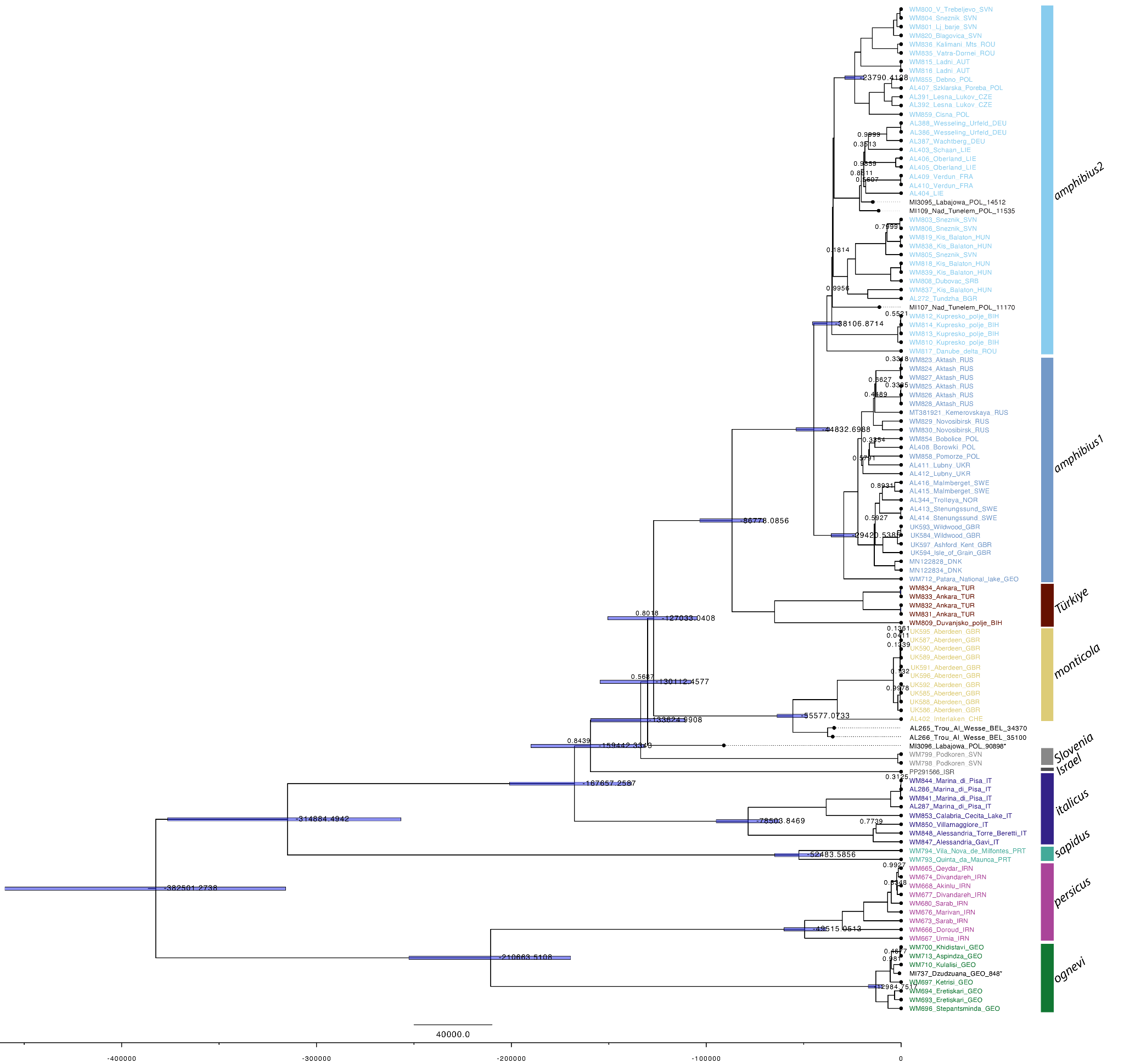

**Figure S1. Tip-calibrated mitochondrial phylogeny of 115 samples of water voles (*Arvicola* sp.) made with BEAST v1.10.4** and visualized in FigTree v.1.4.4. The major clades are highlighted in colour, and split times are displayed with 95% posterior probability. Almost all branches had a posterior probability of 1, and therefore only lower posterior probabilities are shown. Labels represent sample ID, location, and country codes, and for seven ancient samples (black labels) also the age. Two samples highlighted with asterisks (MI3096 and MI737) were dated molecularly, while the others have radiocarbon dates.

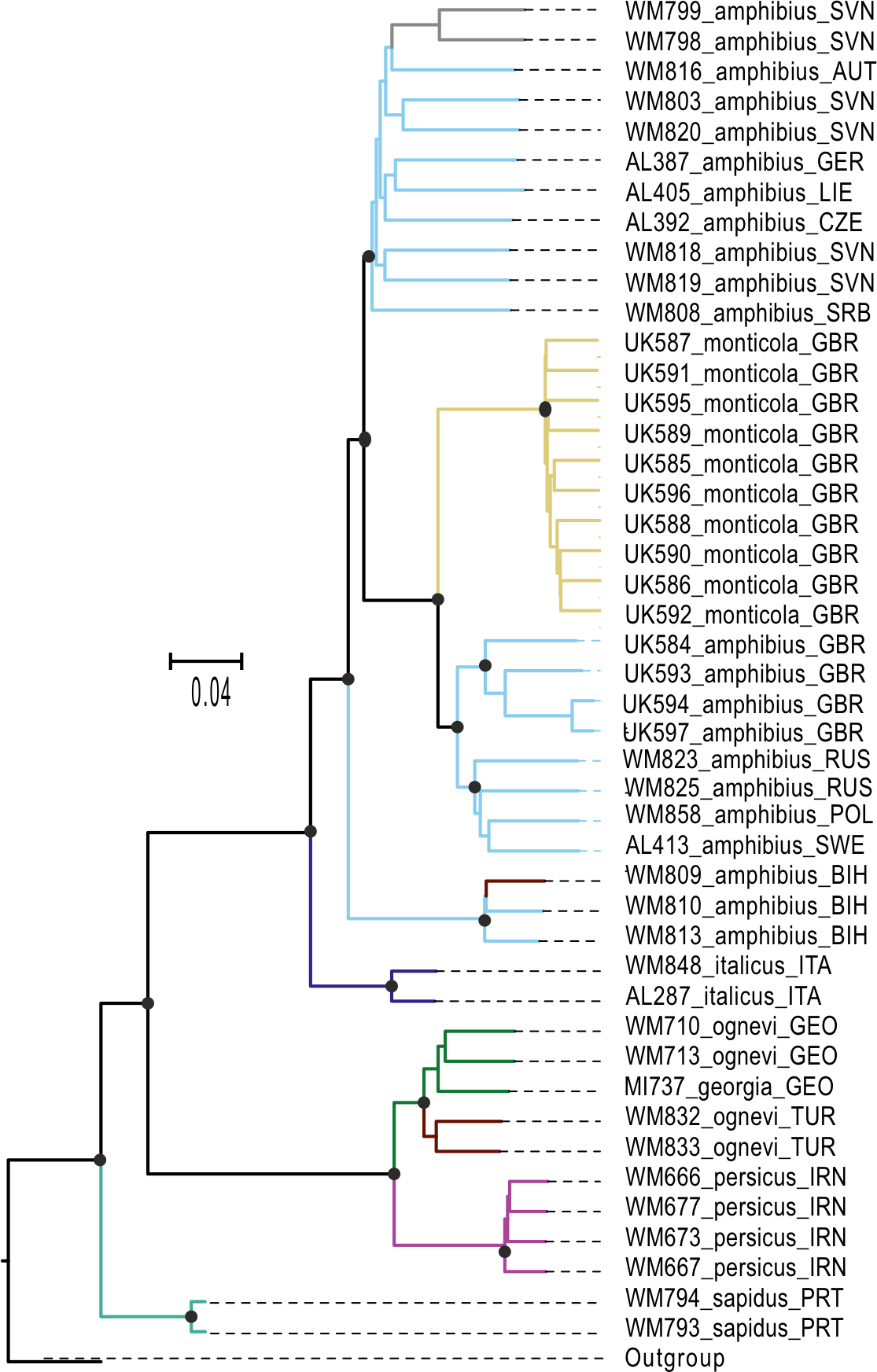

**Figure S2.** **Neighbour-Joining tree of water voles based on identity-by-state distances calculated from nuclear genomic data**

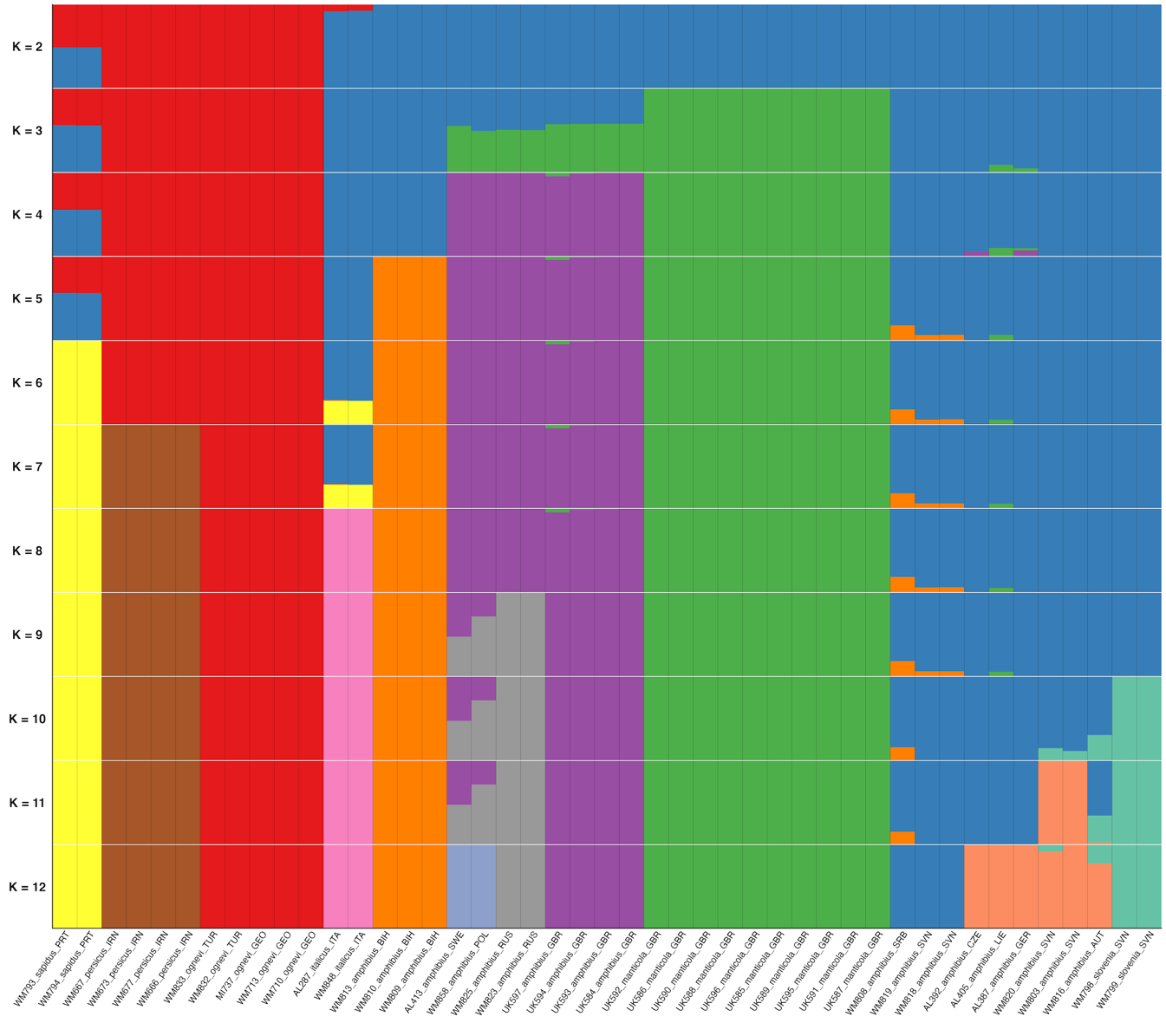
**Fig. S3. Admixture proportions inferred with NGSadmix for K = 2–12 ancestral components.** The analysis was based on 2,307,367 polymorphic sites. Each vertical bar represents an individual, and colours indicate the estimated proportion of ancestry assigned to each inferred ancestral component.

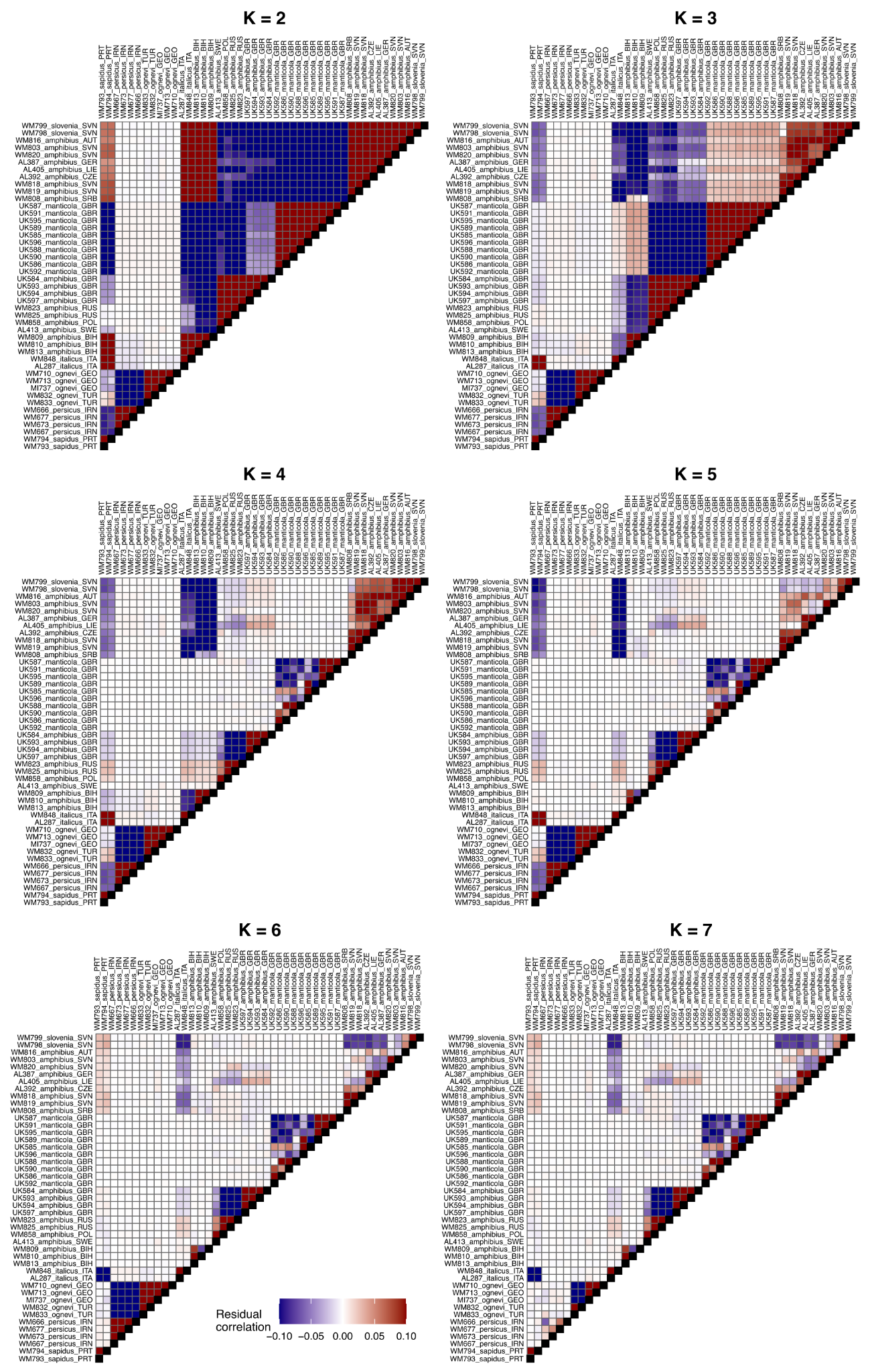

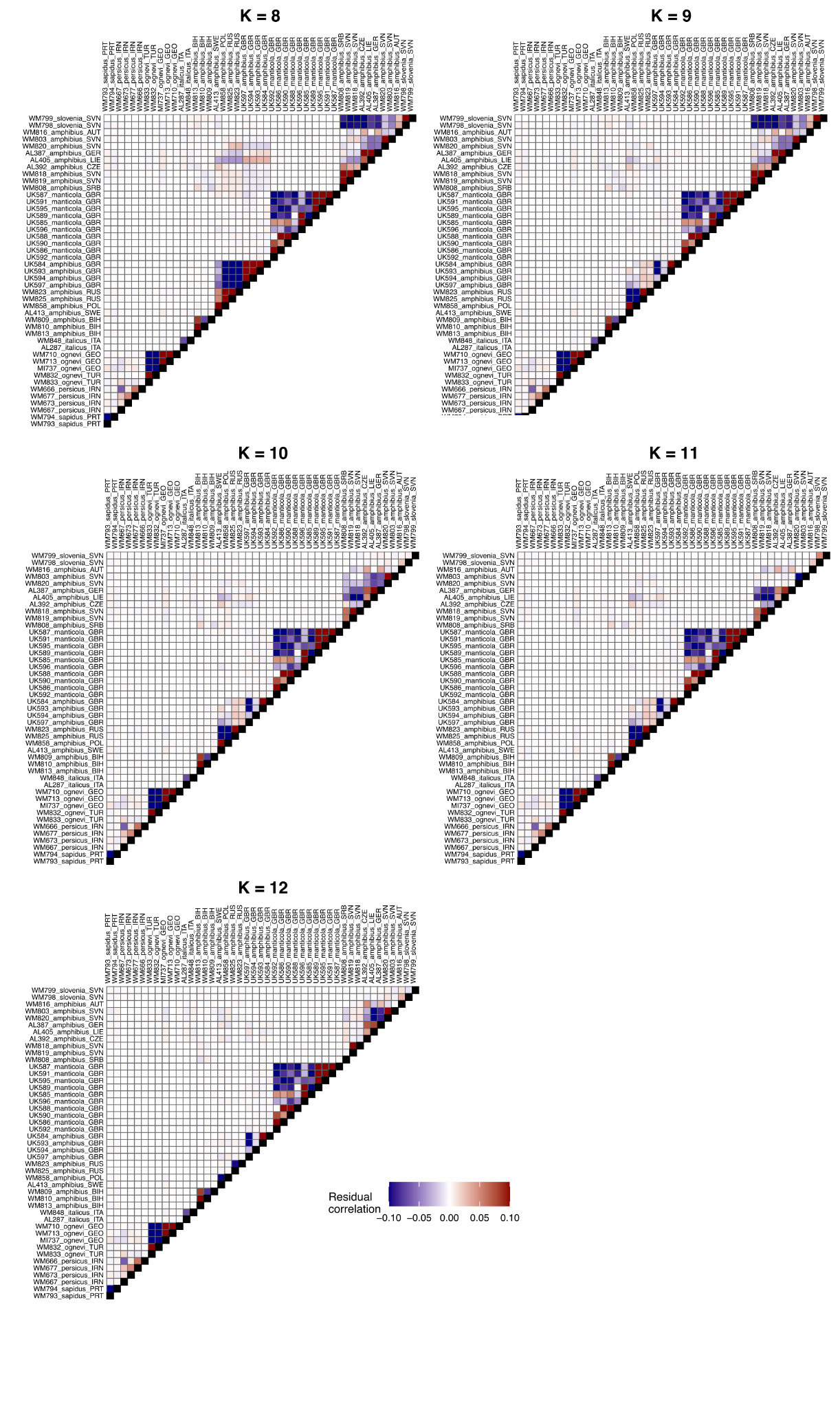

**Figure S4. Evaluation of the estimated admixture proportions using evalAdmix.** (shown in Supplementary Figure S3) using the individual pairwise correlation of residuals estimated with evalAdmix.

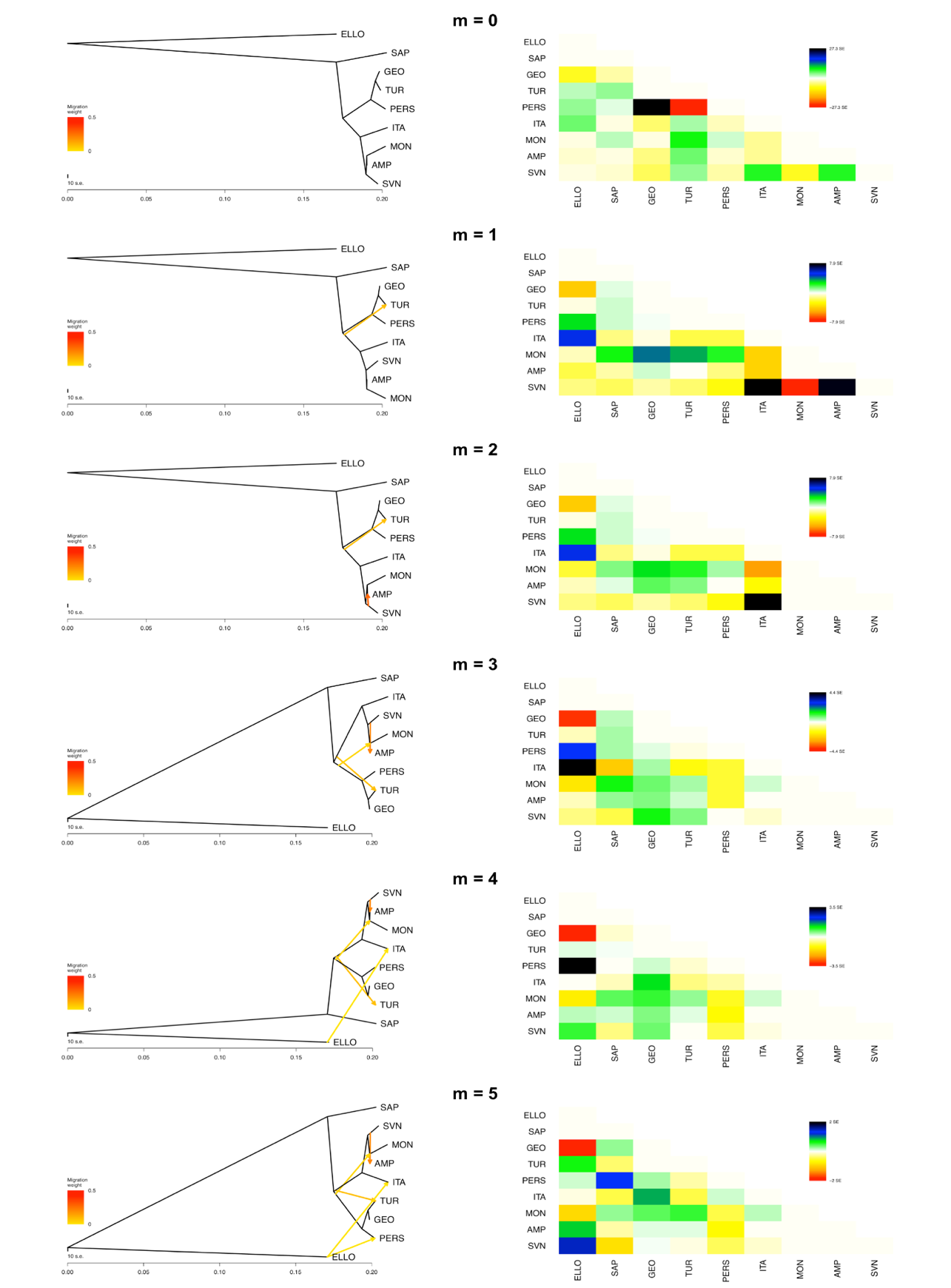

**Figure S5.** **Maximum-likelihood population graphs inferred using TreeMix for water voles divided into eight population groups** (see Table S1 for details), with the number of migration edges ranging from 0 to 5, and the corresponding residual plots.

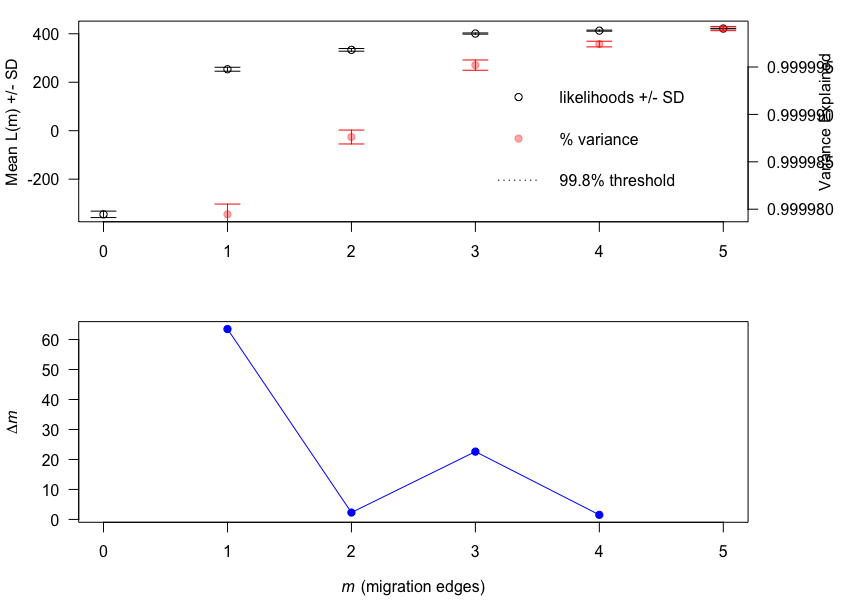

Figure S6. Selection of the number of migration edges in the TreeMix analysis for the dataset partitioned into eight population groups using OptM. The upper panel shows the mean log-likelihood (± SD) across 10 replicate TreeMix runs for models allowing 0–5 migration edges, together with the proportion of variance explained by each model. The dotted line indicates the 99.8% variance-explained threshold. The lower panel shows Δm, the second-order rate of change in likelihood, calculated by OptM to assess the optimal number of migration edges.

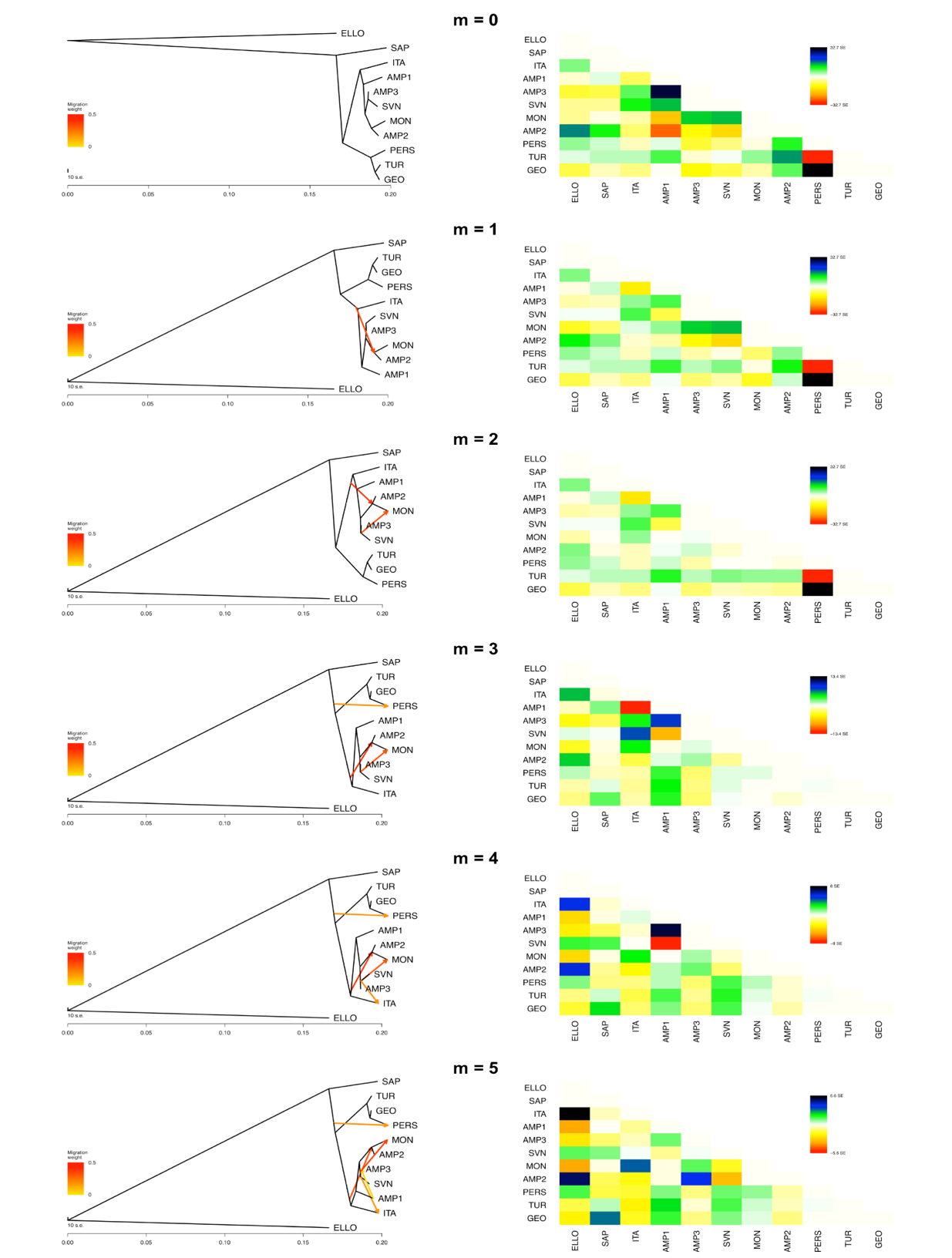
**Figure S7.** **Maximum-likelihood population graphs inferred using TreeMix for water voles divided into ten population groups** (see Table S1 for details), with the number of migration edges ranging from 0 to 5, and the corresponding residual plots.

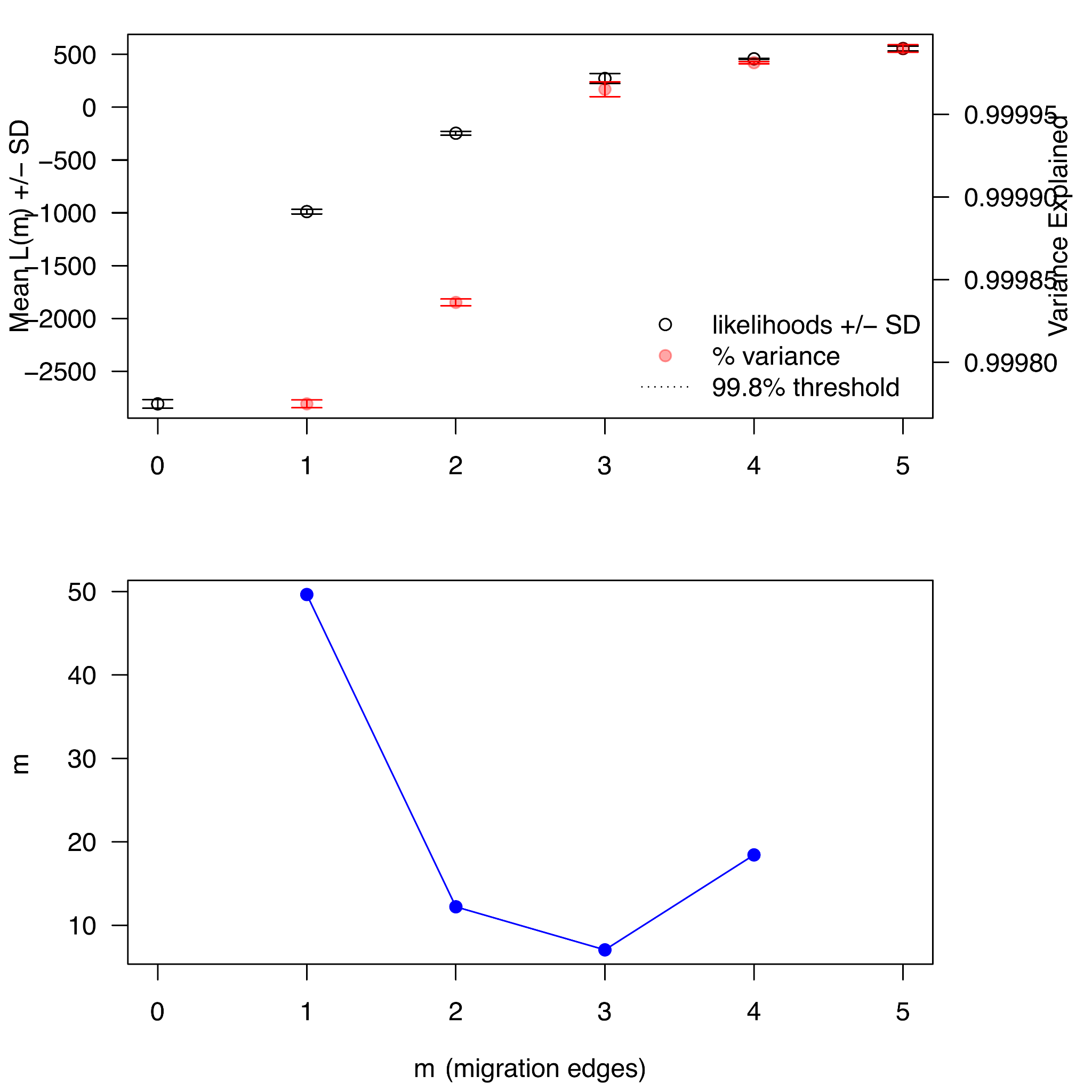

**Figure S8. Selection of the number of migration edges in the TreeMix analysis for the dataset partitioned into eleven population groups using OptM.** The upper panel shows the mean log-likelihood (± SD) across 10 replicate TreeMix runs for models allowing 0–5 migration edges, together with the proportion of variance explained by each model. The dotted line indicates the 99.8% variance-explained threshold. The lower panel shows Δm, the second-order rate of change in likelihood, calculated by OptM to assess the optimal number of migration edges.

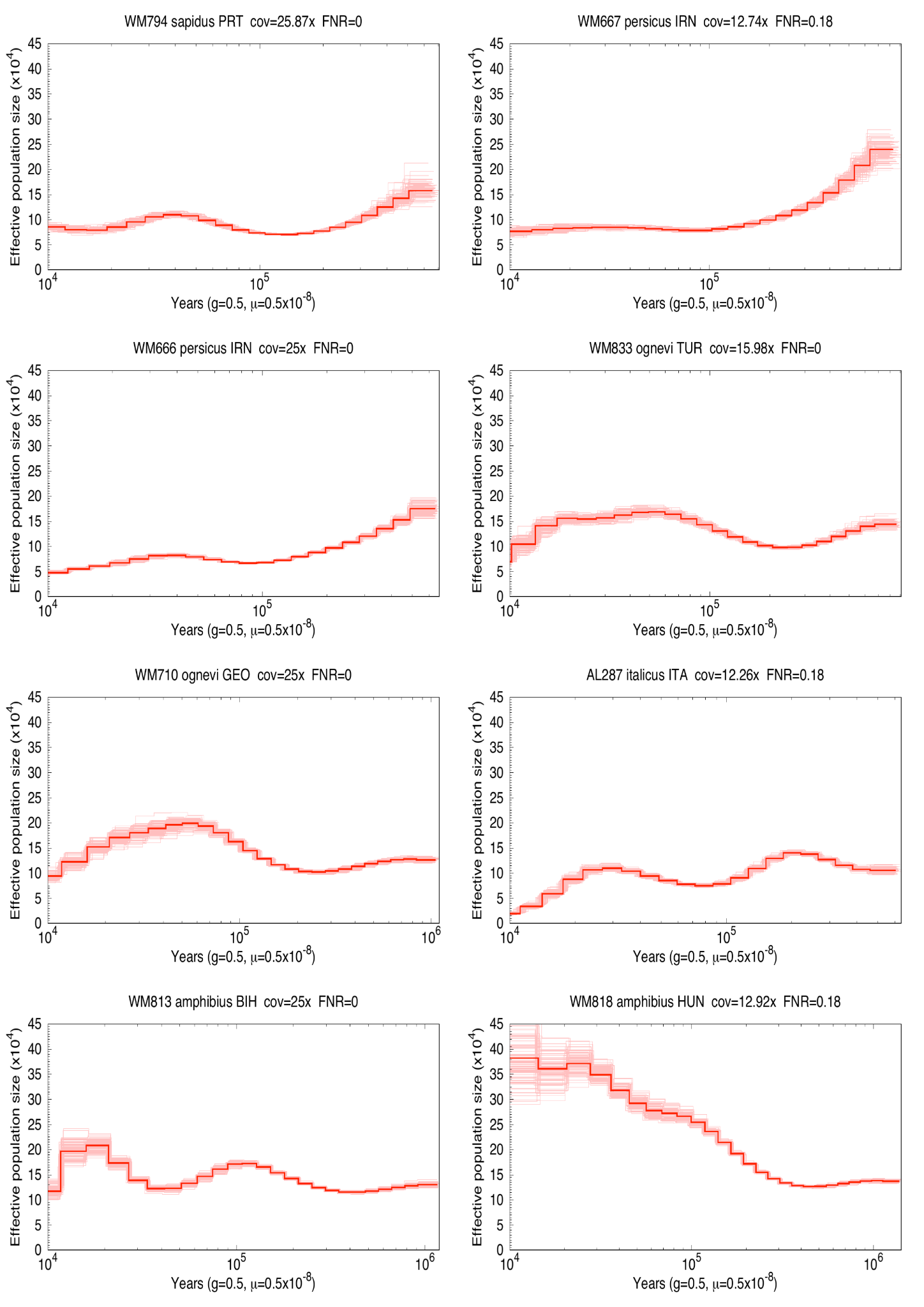

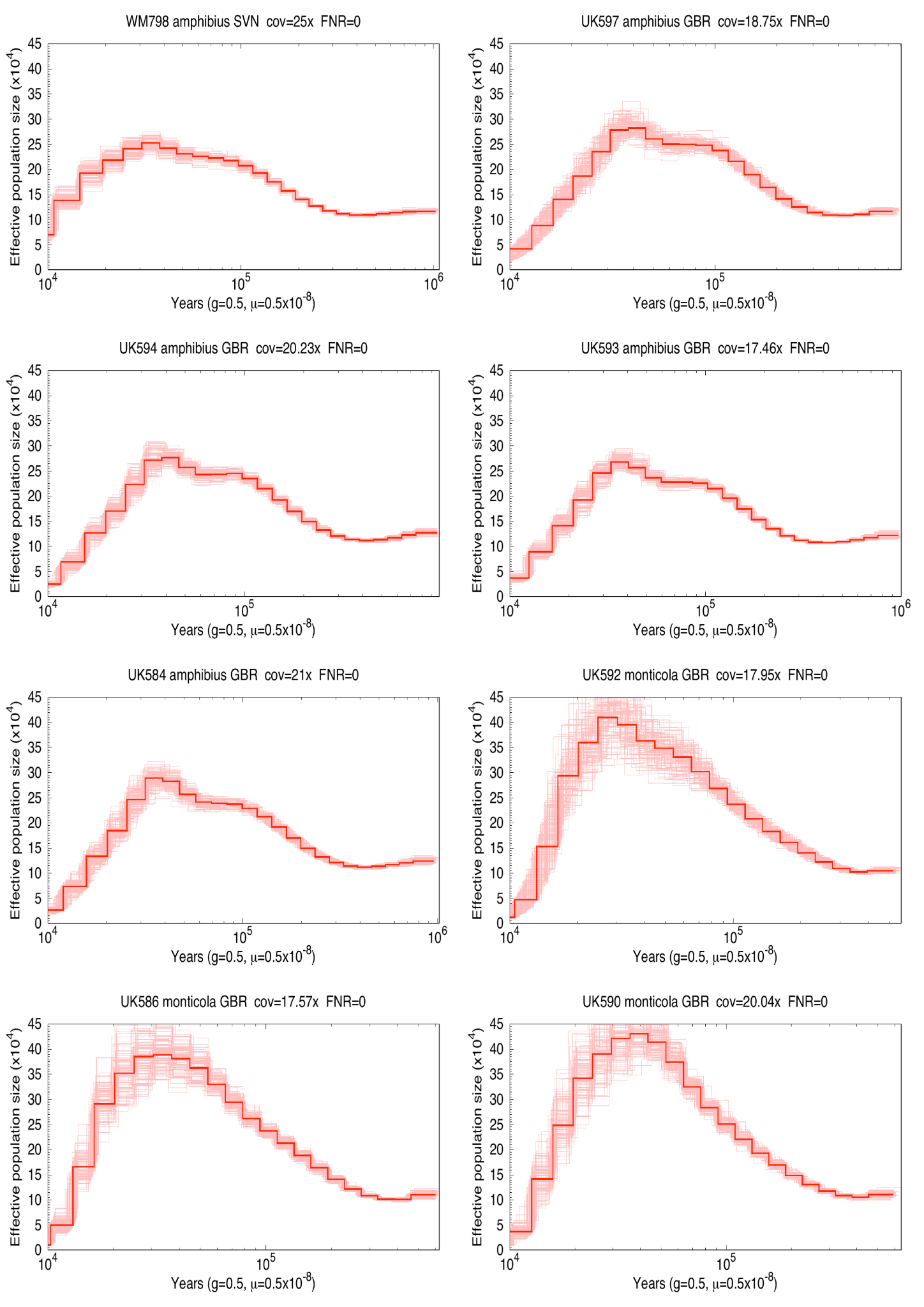

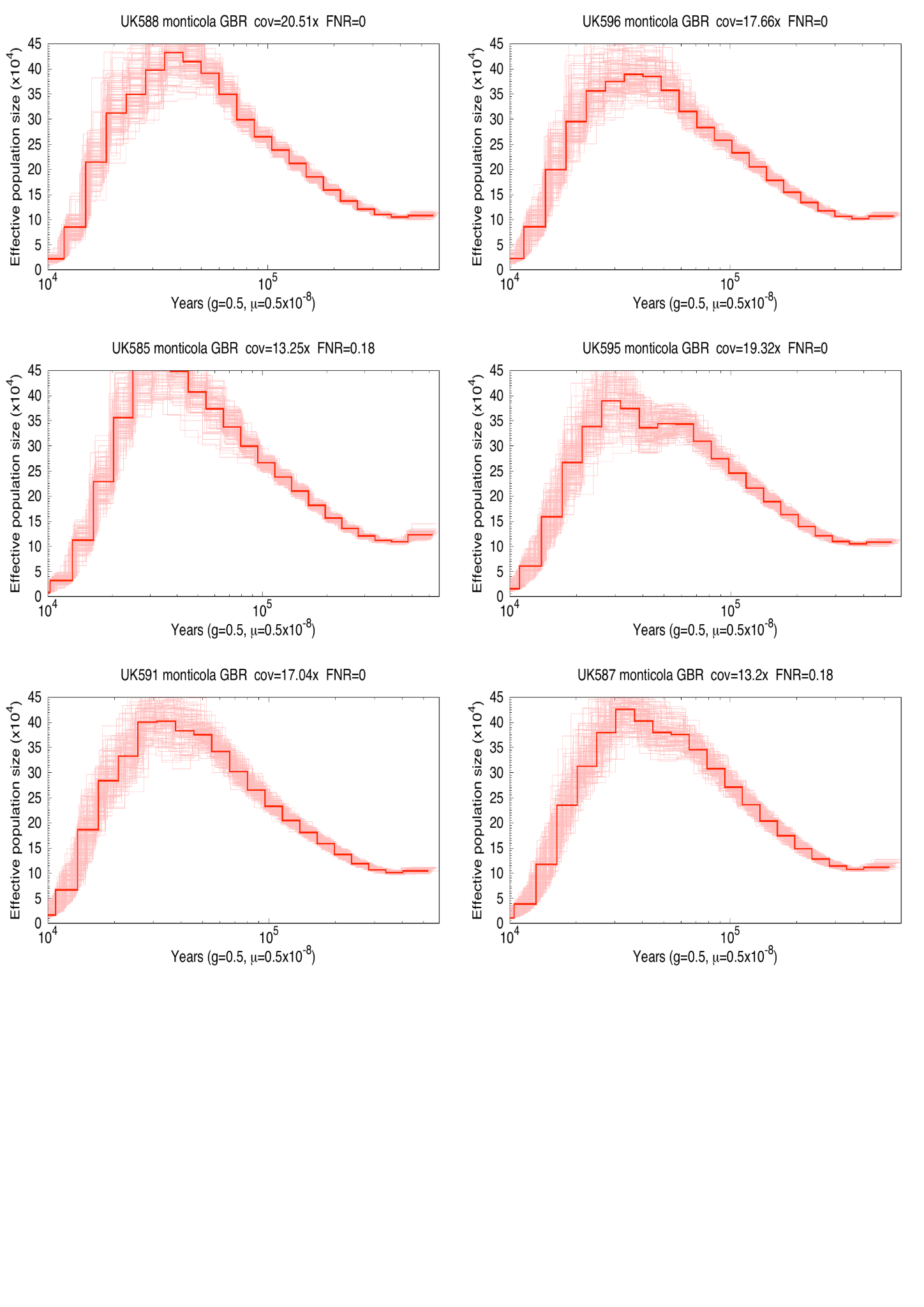

**Figure S9. Demographic reconstruction of water voles using the pairwise sequentially Markovian coalescent (PSMC) model.** A separate plot for each sample with mean genomic coverage above 12× is presented. The PSMC curve was scaled using a mutation rate of 5×10−9 substitutions per site per generation, and assuming two generations per year. The y-axis shows the effective population size and the x-axis shows the time in years on the log scale. Lighter coloured lines represent 100 bootstrap replicates. For samples with a mean genomic coverage below 15×, a false-negative rate (FNR) correction of 0.18 was applied. This correction was estimated by downsampling high-coverage genomes to lower coverage levels and quantifying the resulting loss of heterozygous sites.

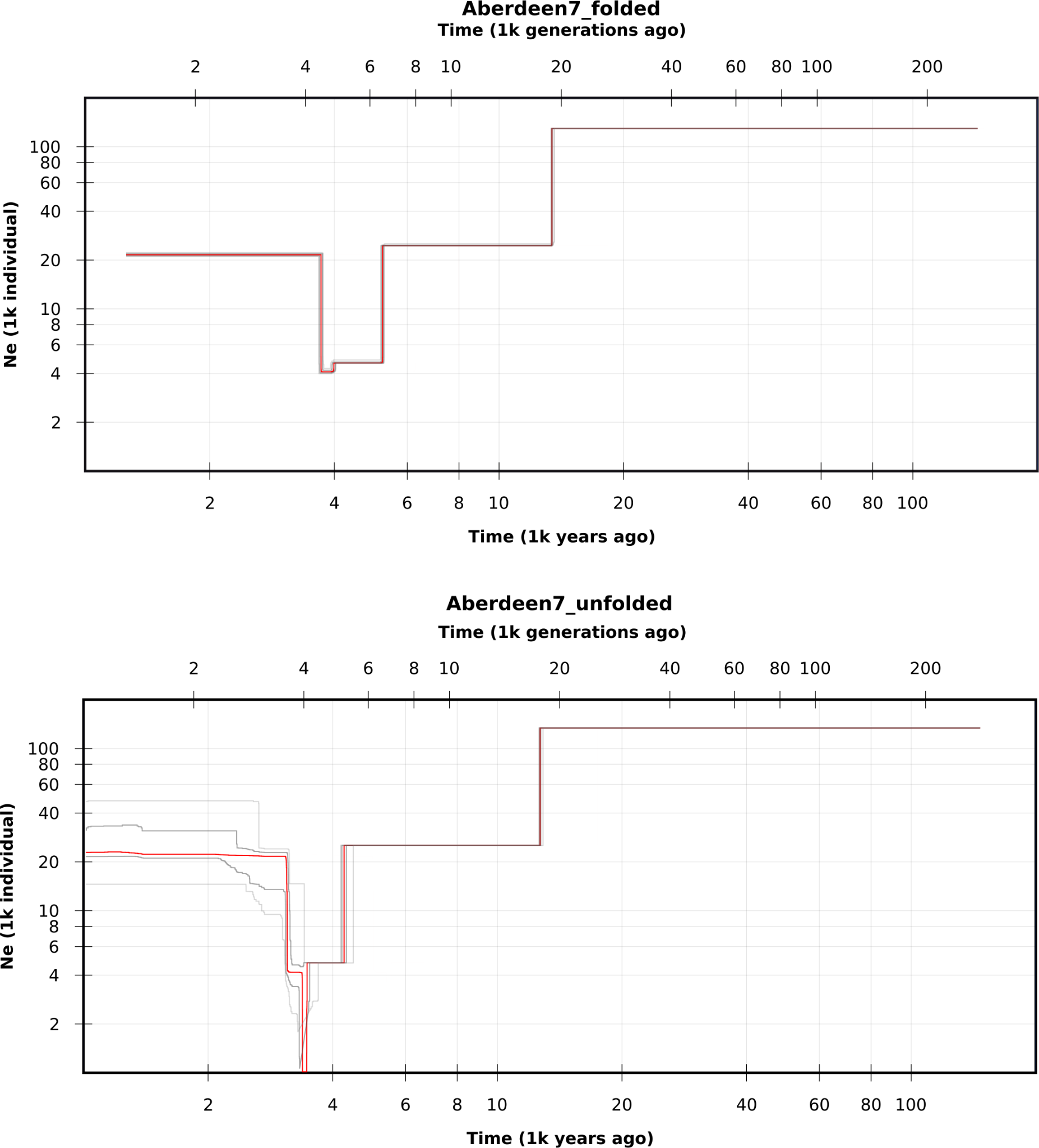

**Figure S10. Demographic history of the Aberdeen water vole population inferred using StairwayPlot2.** The upper panel shows the demographic history inferred from the folded site frequency spectrum, whereas the lower panel shows the corresponding inference based on the unfolded site frequency spectrum. The effective population size (Ne) was scaled using a mutation rate of 5×10−9 substitutions per site per generation, and assuming two generations per year. The y-axis shows the Ne on the log scale. The upper x-axis shows the time in generations and the lower x-axis shows the time in years both on the log scale. The red curve represents the median estimate while grey curves indicate the confidence interval.

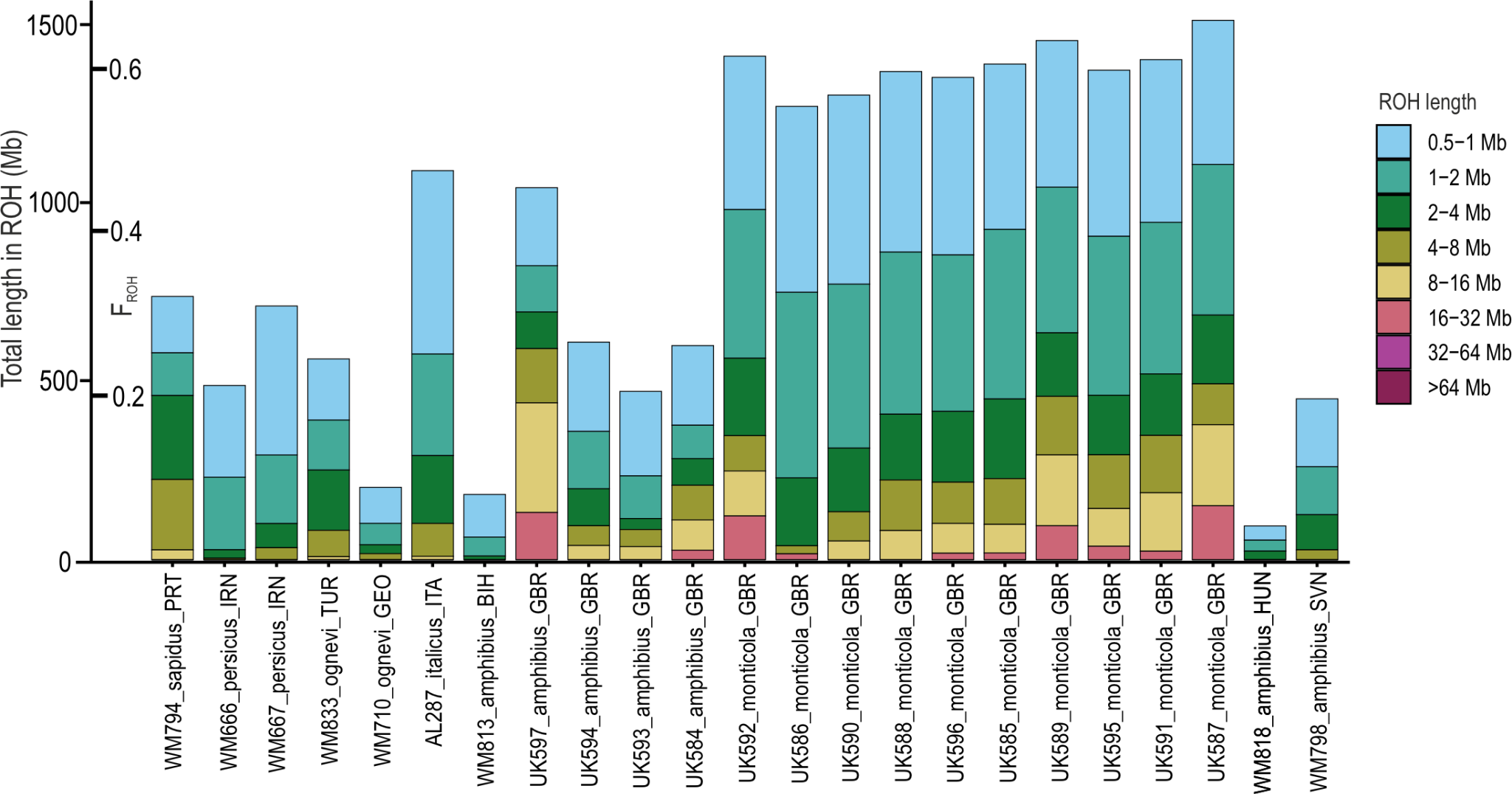

**Figure S11. Runs of homozygosity inferred using PLINK v.1.9 for samples with coverage >12×.** Stacked bars represent the total length of ROHs, partitioned into ROH length classes. The corresponding FROH values were calculated as the summed ROH length divided by the total autosomal genome length (2.3Gb) and are indicated on the secondary scale of the left axis.

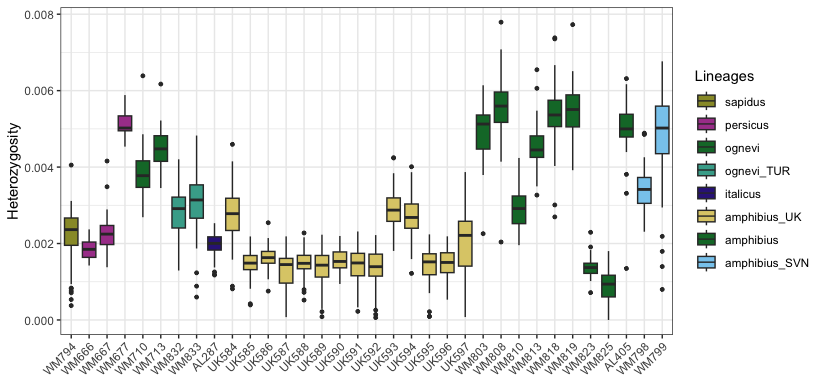

**Figure S12. Genome-wide heterozygosity estimated using ANGSD for samples with mean genomic coverage >5×**. Heterozygosity was calculated in non-overlapping 20 Mb genomic windows. For each individual, boxplots show the median heterozygosity across windows, with boxes representing the interquartile range, whiskers extending to 1.5× the interquartile range, and points indicating values outside this range. Samples are coloured according to mtDNA lineage.

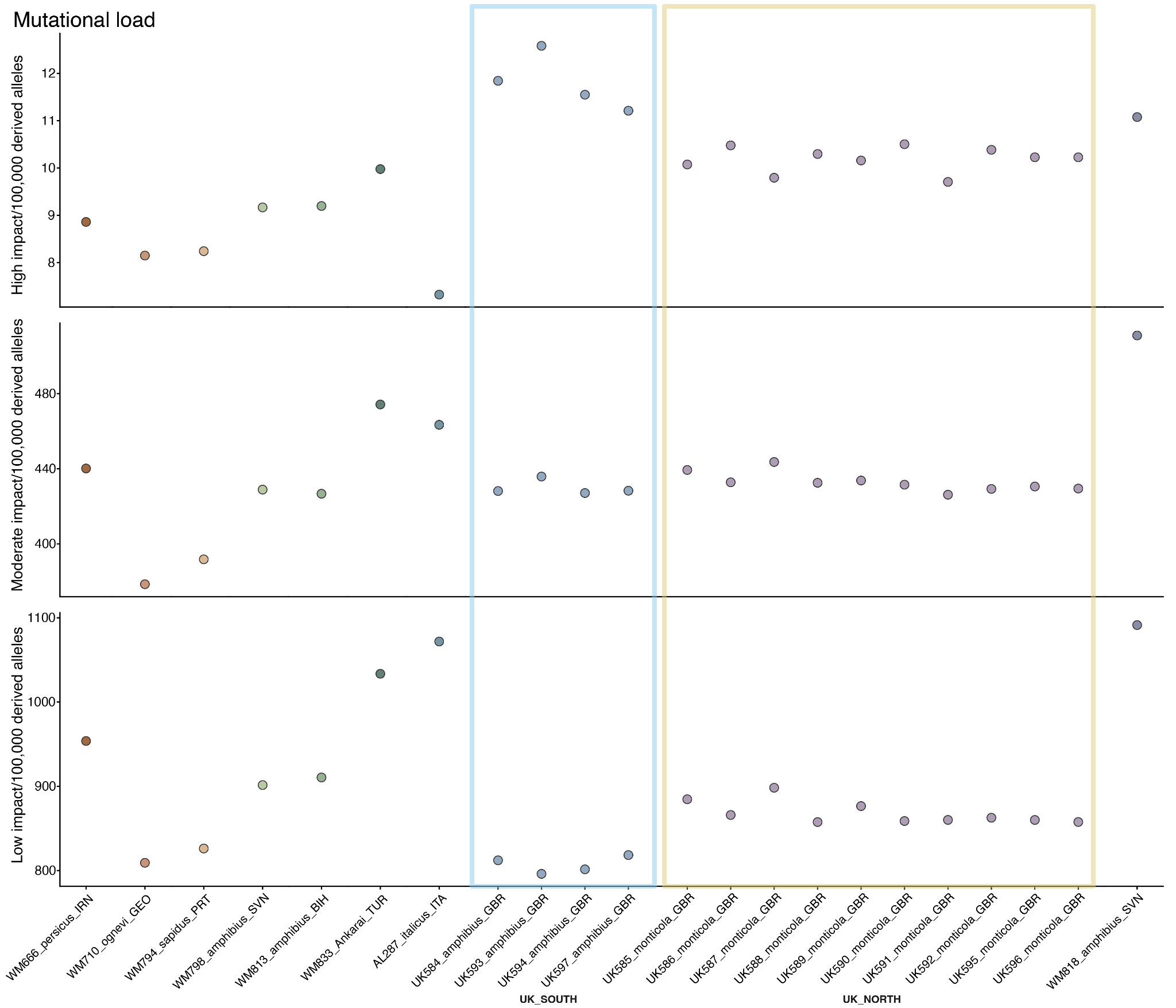

**Figure S13. Mutational load in modern water voles (*Arvicola* sp.).** High, moderate, and low impact mutational load was measured as the number of non-reference (derived) alleles/100,000 derived alleles per individual in each impact class. The blue box highlights samples from southern UK, which belong to the *A. amphibius* lineage, and the yellow box highlights samples from northern UK, which belong to the *A. monticola* lineage.

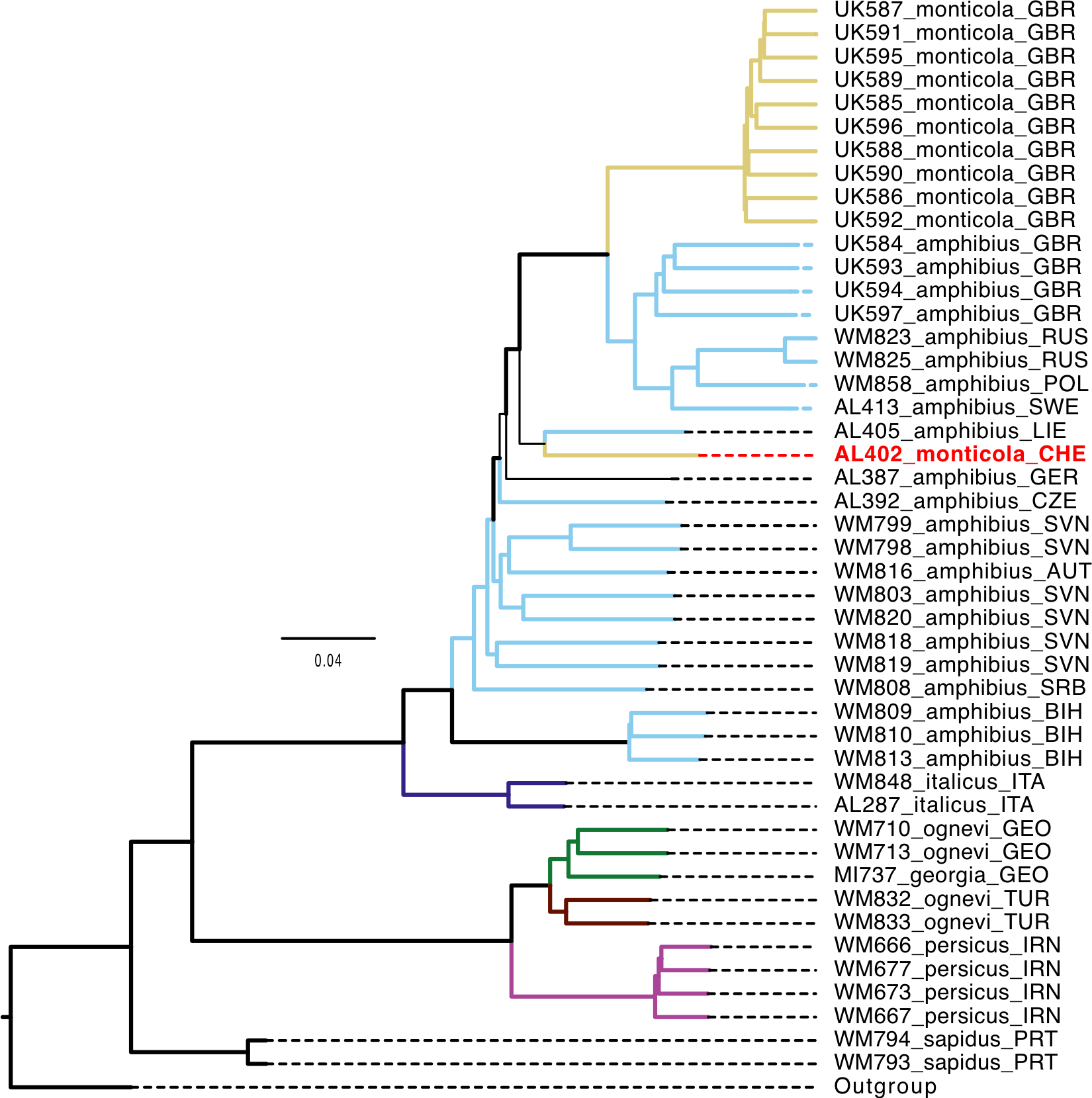

**Figure S14. Neighbour-Joining tree of water voles based on identity-by-state distances calculated from nuclear genomic data.** The tree includes the low coverage (0.04×) sample AL402.

### 2 Additional Methods and Results

#### 2.1 DNA extraction, sequencing and data processing

Prior to DNA extraction, modern dry tissue samples were incubated in water for 3h and fragmented into 2 mm diameter pieces, while modern bone samples labelled “WM” were pre-treated by incubation in 500 µl 0.5% hypochlorite, rinsed twice in ultrapure water, and crushed with a pipette tip into smaller pieces. After extraction, quality checks were performed including measurements on DeNovix NanoDrop spectrophotometer and 1% agarose gel, and the extracts were sonicated to ca. 300bp using Covaris and size selected using a 0.6x ratio of magnetic beads. The double-stranded and double-indexed libraries were amplified using AmpliTaq Gold 360 Master Mix (Applied Biosystems), and the number of cycles required for sufficient amplification was assessed by real-time chain reaction (qPCR). For AL-labelled specimens, amplification of double-stranded libraries was performed using AccuPrime Pfx DNA polymerase (Thermo Fisher Scientific), while amplification of single-stranded libraries was performed with AmpliTaq Gold DNA polymerase (Applied Biosystems), and qPCR was used to assess the required number of cycles. The indexed libraries were pooled together, and size-selected for using Sera-Mag™ beads (Cytiva) prior to sequencing. One indexed library per sample was prepared for initial low depth sequencing, and for deep sequencing additional indexed libraries were added; 15 for AL287, eight for AL392, seven for AL387, five for AL405, and four for AL413, to maintain complexity.

The raw fastq-files generated for the modern AL-labelled samples were trimmed and merged with Fastp v.0.24.0 (Chen et al. 2018), and only merged reads with a minimum length of 30 bp were kept. After competitive mapping, short reads (<70 bp) were mapped with bwa aln with default parameters for ancient aDNA (−o 2 −n 0.01 −l 1024), while longer reads were mapped with bwa mem using default parameters. Only bam-files with reads mapping to the target reference were used for further analyses. Reads with mapping quality <25 were discarded, BAM-files were merged per library and duplicates removed. Libraries from the same sample sequenced with different indexes and/or on different lanes were merged and duplicates removed once more.

#### 2.2 mtDNA phylogeny

We included 115 mitochondrial genomes in the phylogeny; 97 that were generated for the purpose of this study, including 90 modern and seven ancient, and 18 published modern genomes. Six of the ancient samples were radiocarbon dated, but MI3096 was estimated to have an infinite radiocarbon date, and MI737 had no radiocarbon date and both were therefore included for molecular dating. The sequences were aligned using MAFFT v.7.526 (Katoh and Standley 2013), and the alignment was visually inspected. The ends were trimmed to ensure that all mitogenomes were of equal lengths using a custom python script, to a final length of 16054 bp. PartitionFinder2 (Lanfear et al. 2016) was used to identify the most suitable partitioning scheme and substitution model (Table S3). BEAST v.1.10.4 was used to construct the mitochondrial phylogeny, molecularly date the two ancient samples, and estimate divergence times between the major lineages. We used eight partitions for all analyses based on the results from PartitionFinder2, with unlinked substitution models but the same clock model applied to all partitions. We then used Bayesian evaluation of the temporal signal (BETS) to test if our dataset contained a temporal signal strong enough to calibrate the molecular clock. We compared four models; (I) with tip dating and a strict clock, (II) without tip dating and a strict clock, (III) with tip dating and an uncorrelated relaxed lognormal clock (UCLN), and (IV) without tip dating and UCLN clock. A coalescent constant population size tree prior was applied to all models, and three independent runs running for 100 M generations each and sampled every 10,000 steps were generated. The log marginal likelihood (MLE) was estimated using generalized stepping-stone sampling with 100 stepping stones and 1 million iterations. All runs were inspected in Tracer v.1.7.2 for convergence, and ESS was >200 for all parameters. The mean values from the MLE calculations were calculated and then compared, resulting in strong support for a temporal signal (e^x^(6,933)=1025,566), and a strict clock model (e^x^(4,739)=114,320) (Table S5). In addition, we evaluated which tree prior was the best fit by comparing model (V): tip dating, strict clock, coalescent: Bayesian SkyGrid, to model (I): tip dating, strict clock, coalescent: constant size, which generated support for the Bayesian SkyGrid tree prior (e^x^(1,317)=3,732). The three tree-files generated from model V were summarized using LogCombiner v.1.10.4, with 10% burn-in and a resample frequency of 20,000. TreeAnnotator v1.10.4 was used to create a consensus tree based on 13,501 trees, and the final tree was visualized in FigTree v1.4.4 (Rambaut 2018).

**Table S5.** Results of Bayesian evaluation of the temporal signal and tree prior.

| ID | Description | logML | mean logML |
| --- | --- | --- | --- |
| Model I | Tips; strict clock; coalescent constant size tree prior | -61219,360 | -61216,210 |
|  |  | -61214,600 |  |
|  |  | -61214,669 |  |
| Model II | No tips; strict clock; coalescent constant size tree prior | -62241,266 | -62241,838 |
|  |  | -62243,345 |  |
|  |  | -62240,903 |  |
| Model III | Tips; UCLN clock; coalescent constant size tree prior | -61329,414 | -61330,531 |
|  |  | -61329,254 |  |
|  |  | -61332,926 |  |
| Model IV | No tips; UCLN clock; coalescent constant size tree prior | -5,113E69 | Failed |
|  |  | -7,343E83 |  |
|  |  | -1,453E82 |  |
| Model V | Tips; strict clock; coalescent Bayesian SkyGrid tree prior | -61217,814 | -61219,943 |
|  |  | -61217,814 |  |
|  |  | -61224,202 |  |

#### 2.3 Nuclear DNA Neighbour-Joining phylogeny

To assess node support in the NJ phylogeny reconstructed from IBS distances, we generated 1,000 bootstrap replicates by resampling the IBS output file produced by ANGSD with replacement using 50-kb genomic blocks and recalculating pairwise IBS distances for each replicate. Resampling was performed using a custom Python script Block_resample_IBS1_file_v2.py (<https://github.com/mateuszbaca/Arvicola-commands/blob/main/Full_dataset/01_GL_and_IBS/Block_resample_IBS1_file_v2.py>). The NJ tree for each bootstrap distance matrix was computed using RapidNJ (Simonsen et al. 2008). Node support was calculated as the proportion of bootstrap trees recovering each clade using *ape* package in R.

#### 2.4 ngsRelate

Pairwise relatedness among individuals from the northern UK (Aberdeen) water vole population was assessed using ngsRelate (Korneliussen and Moltke 2015), which estimates relatedness directly from genotype likelihoods. Genotype likelihoods were generated with ANGSD using the same filtering criteria as for the population-genomic analyses. The analysis identified four pairs of individuals showing levels of relatedness consistent with second-degree relationships (Table S6). To minimise the influence of related individuals on demographic inference, three individuals (with the lowest genomic coverage were removed UK585, UK587, UK591), leaving seven unrelated individuals for downstream StairwayPlot2 analyses.

**Table S6.** Pairwise relatedness estimates for the northern UK (Aberdeen) water voles identified as potential second-degree relatives. rab is the pairwise relatedness coefficient estimated by ngsRelate, θ is the estimated kinship coefficient. R0 and R1 represent the estimated probabilities of sharing zero and one allele identical by descent, respectively; KING denotes the KING-robust kinship coefficient. Four pairs showed kinship coefficients consistent with second-degree relatedness.

| **Sample 1** | **Sample 2** | **rab** | **θ** | **R0** | **R1** | **KING** | **Potential relationship** |
| --- | --- | --- | --- | --- | --- | --- | --- |
| UK587 | UK591 | 0.27786 | 0.13893 | 0.09678 | 0.61738 | 0.22891 | 2nd degree |
| UK591 | UK595 | 0.26050 | 0.13025 | 0.06562 | 0.62433 | 0.24568 | 2nd degree |
| UK587 | UK595 | 0.22764 | 0.11382 | 0.11438 | 0.60026 | 0.21715 | 2nd degree |
| UK585 | UK596 | 0.17059 | 0.08530 | 0.12924 | 0.46448 | 0.18429 | 2nd–3rd degree |

#### 2.5 StairwayPlot2

Historical changes in effective population size were estimated using Stairway Plot 2. Analyses were performed using both folded and unfolded site frequency spectra obtained from seven unrelated individuals from the Aberdeen population. Site allele-frequency likelihoods were generated in ANGSD v. 0.930 from the filtered set of genomic positions specified with -sites using the following parameters: -uniqueOnly 1, -remove_bads 1, -minMapQ 25, -minQ 20, -doCounts 1, -GL 2, -doSaf 1, -doMaf 1, -doMajorMinor 1, -minInd 6, -P 6 and converted to SFS using realSFS. StairwayPlot v2.2 was run with 14 sampled chromosomes, 67% of sites used for training, random breakpoint numbers of 3, 6, 9 and 12, and 200 input replicates. Demographic histories were scaled assuming a mutation rate of 5×10^−9^ substitutions per site per generation and a generation time of 0.5 years.

#### 2.6 TreeMix

To generate TreeMix input files, samples were divided into either 11 or 8 groups including *Ellobius talpinus* as an outgroup. For each group, site allele-frequency likelihoods (SAFs) were estimated using ANGSD v0.930 at a filtered set of genomic positions specified with -sites, applying the following parameters: -uniqueOnly 1, -remove_bads 1, -minMapQ 25, -minQ 20, -doCounts 1, -GL 2, -doSaf 1, -doMaf 1, and -doMajorMinor 1, with the ArvAmp1.2 reference allele treated as ancestral. Group-specific SAF files were combined using realSFS print, retaining positions with data across all groups. A custom Python script (<https://github.com/mateuszbaca/Arvicola-commands/blob/main/Full_dataset/saf2treemix3.1.py>) was then used to identify the most likely derived allele count for each group at each position, based on the maximum SAF likelihood. These counts were converted into ancestral and derived allele counts in TreeMix format, excluding sites fixed for the same allele across all groups. This produced a file containing 149,146,928 positions. To reduce linkage among sites and computational burden the file was thinned thirty times to 5,100,930 positions using a custom Python script. Then 10 replicates for each number of migration edges from 0 to 5 were run using the parameters as follows: -global, -m, n_warn 1, -root, -seed. Then the replicate with the highest likelihood was selected for visualization. In parallel, to enable the use of *optM*, we performed 10 replicate runs for each number of migration edges, each using a randomly selected subset comprising 70% of the positions in the full dataset, to introduce variation in the resulting model likelihoods.

#### 2.7 FastSimCoal2

To evaluate whether gene flow between Georgian Arvicola ognevi and A. persicus could affect estimates of their divergence time obtained with the Two-Two (TT) method, we performed demographic modelling using fastsimcoal2 v2.8. The analyses were based on the joint two-dimensional site-frequency spectrum (2D-SFS) calculated from three Georgian ognevi and four Iranian A. persicus genomes. We used population-specific SAFs estimated using ANGSD for the TreeMix analysis (Supplementary Text 2.6). The joint SFS was estimated with realSFS print. In the final fastsimcoal2 analyses, invariant sites were retained. Four alternative demographic scenarios were compared: (i) **No gene flow**, in which the two populations diverged from a common ancestral population and evolved in complete isolation; (ii) **Early gene flow**, allowing continuous migration between the populations during an early period following divergence but no recent migration; (iii) **Pulse**, a single pulse of gene flow between the populations; and (iv) **Recent gene flow**, representing secondary contact, with a period of isolation following divergence and continuous bidirectional migration from an estimated time in the past until the present.

Population divergence time (TDIV), ancestral and descendant effective population sizes and, where applicable, migration rates and the timing of changes in gene flow (TCHNG) were estimated by composite-likelihood optimization. We used identical, population-size bounds derived from the levels of genomic heterozygosity observed in the two analysed populations. The ancestral effective population size and one descendant population size were allowed to vary between 100,000 and 500,000 individuals, whereas the second descendant population size was allowed to vary between 50,000 and 500,000 individuals; log-uniform priors were used in all cases. Divergence time and gene-flow parameters were otherwise left free to vary within the default parameter ranges (Table S7). For each likelihood evaluation, 100,000 coalescent simulations were performed, and parameter optimization was carried out for 40 ECM cycles. SFS entries with fewer than 10 observations were excluded using −C 10. At least 30 independent optimization runs were performed for each demographic model.

**Table S7.** Parameters and search ranges of four demographic models analysed with *fastsimcoal2.* NANC: ancestral effective population size; NMR: Georgian *ognevi* effective population size; NMB: *A. persicus* effective population size; TDIV: population split time (generations); TCHNG/TCHANG: timing of change in gene flow (generations); TPulse: timing of pulse admixture; MIG1 and MIG2: migration parameters as defined in the model templates. Estimates are from the highest composite-likelihood run under the standardized N_e_ constraints.

| **No** | **Model** | **Parameter** | **Value Type** | **Distribution Type** | **Minimum** | **Maximum** | **Bounded** | **Estimate** |
| --- | --- | --- | --- | --- | --- | --- | --- | --- |
| **1** | **Recent gene flow** | NANC | Integer | log-uniform | 100000 | 500000 | No | 125402 |
|  |  | NMR | Integer | log-uniform | 100000 | 500000 | No | 121108 |
|  |  | NMB | Integer | log-uniform | 50000 | 500000 | No | 55838 |
|  |  | TDIV | Integer | log-uniform | 50000 | 2000000 | No | 812354 |
|  |  | TCHNG | Integer | log-uniform | 1000 | TDIV | Yes | 634284 |
|  |  | MIG1 | Float | log-uniform | 0.00 | 0.01 | No | 0.0000287 |
|  |  | MIG2 | Float | log-uniform | 0.00 | 0.01 | No | 0.0000049 |
| **2** | **Early gene flow** | NANC | Integer | log-uniform | 100000 | 500000 | No | 416266 |
|  |  | NMR | Integer | log-uniform | 100000 | 500000 | No | 426413 |
|  |  | NMB | Integer | log-uniform | 50000 | 500000 | No | 438088 |
|  |  | TDIV | Integer | log-uniform | 50000 | 2000000 | No | 1620418 |
|  |  | TCHANG | Integer | log-uniform | 1000 | TDIV | Yes | 1156 |
|  |  | MIG1 | Float | log-uniform | 0.00 | 0.01 | No | 0.0000013 |
|  |  | MIG2 | Float | log-uniform | 0.00 | 0.01 | No | 0.0000012 |
| **3** | **Pulse gene flow** | NANC | Integer | log-uniform | 100000 | 500000 | No | 453393 |
|  |  | NMR | Integer | log-uniform | 100000 | 500000 | No | 475578 |
|  |  | NMB | Integer | log-uniform | 50000 | 500000 | No | 406007 |
|  |  | TDIV | Integer | log-uniform | 50000 | 2000000 | No | 1708960 |
|  |  | TPulse | Integer | log-uniform | 100 | TDIV | Yes | 46766,0 |
|  |  | Pulse proportion | Float | fixed | 0.01 | 0.01 | Fixed | 0,1 |
| **4** | **No gene flow** | NANC | Integer | log-uniform | 100000 | 500000 | No | 496703 |
|  |  | NMR | Integer | log-uniform | 100000 | 500000 | No | 401020 |
|  |  | NMB | Integer | log-uniform | 50000 | 500000 | No | 428687 |
|  |  | TDIV | Integer | log-uniform | 50000 | 2000000 | No | 236495 |

Models were ranked according to their maximum composite likelihood and Akaike information criterion, (AIC=2k−2\lnL), where (k) denotes the number of free parameters. Times in generations were converted to years assuming a generation time of 0.5 years (Table S8).

The best-fitting Recent-gene-flow model showed a multimodal likelihood surface, with independent optimization runs converging on several distinct high-likelihood combinations of parameters. To improve exploration of the parameter space and reduce the risk of retaining a local optimum, the number of independent optimizations for this model was therefore increased to 1,000.

**Table S8.** Comparison of the four fastsimcoal2 demographic models. AIC = 2k − 2lnL; ΔAIC is relative to the best-fitting model.

| No | **Model** | **Free parameters (k)** | **Best MaxEstLhood** | **AIC** | **ΔAIC** | **Best TDIV (generations)** | **TDIV (ka)** |
| --- | --- | --- | --- | --- | --- | --- | --- |
| **1** | **Recent gene flow** | 7 | -564570507 | 1129141028 | 0.0 | 812354 | 406.2 |
| **2** | **Early gene flow** | 7 | -575208275 | 1150416563 | 21275535.2 | 1620418 | 810.2 |
| **3** | **Pulse gene flow** | 5 | -580369560 | 1160739130 | 31598102.1 | 1708960 | 854.5 |
| **4** | **No gene flow** | 4 | -583738625 | 1167477259 | 38336230.9 | 236495 | 118.2 |

#### 2.8 Runs of homozygosity

Runs of homozygosity were inferred using two complementary approaches: PLINK v.1.9 for samples with mean genomic coverage >12× and ROHan for samples with mean genomic coverage >5×.

For the medium-higher-coverage dataset, ROHs were inferred using a minimum ROH length of 500 kb (--homozyg-kb 500) and at least 150 SNPs per segment (--homozyg-snp 150), a maximum marker density of 10 kb/SNP (--homozyg-density 10), a maximum gap of 500 kb between consecutive SNPs (--homozyg-gap 500), and a 50-SNP sliding window (--homozyg-window-snp 50) allowing one heterozygous genotype (--homozyg-window-het 1) and twenty missing genotypes (--homozyg-window-missing 20); SNPs were required to occur in at least 5% of homozygous windows (--homozyg-window-threshold 0.05), and up to three heterozygous genotypes were permitted per ROH (--homozyg-het 3).

We tested the impact of the number of missing genotypes allowed per sliding window (--homozyg-window-missing; HWM) on ROH discovery by repeating the analysis with thresholds of 5, 10, 15, 20, and 25 missing genotypes per 50-SNP window, while keeping all other parameters unchanged (Table S9). Higher-coverage samples (generally >~15×) were largely insensitive to this setting, whereas several lower-coverage samples showed substantial increases in the total inferred ROH length as HWM was relaxed. In the most affected samples, total ROH length increased steeply with increasing HWM values and generally approached a plateau between HWM20 and HWM25 (Figure S15). We therefore used --homozyg-window-missing 20 in the final analyses.

**Table S9.** Total length of identified ROH segments (Mb) depending on the maximum number of missing genotypes per window.

| Sample | mean genomic coverage | HWM5 | HWM10 | HWM15 | HWM20 | HWM25 |
| --- | --- | --- | --- | --- | --- | --- |
| AL287_italicus_ITA | 12,26 | 0 | 76 | 684 | 1085 | 1154 |
| WM667_persicus_IRN | 12,74 | 590 | 594 | 595 | 597 | 598 |
| WM818_amphibius_HUN | 12,92 | 24 | 780 | 1323 | 1382 | 1389 |
| UK587_monticola_GBR | 13,2 | 1157 | 1255 | 1261 | 1264 | 1265 |
| UK585_monticola_GBR | 13,25 | 11 | 634 | 1395 | 1504 | 1513 |
| UK589_monticola_GBR | 14,48 | 1338 | 1355 | 1359 | 1361 | 1362 |
| WM833_ognevi_TUR | 15,98 | 290 | 1326 | 1434 | 1447 | 1449 |
| UK591_monticola_GBR | 17,04 | 1274 | 1292 | 1295 | 1295 | 1296 |
| UK593_amphibius_GBR | 17,46 | 1288 | 1388 | 1392 | 1394 | 1395 |
| UK586_monticola_GBR | 17,57 | 1371 | 1398 | 1401 | 1404 | 1406 |
| UK596_monticola_GBR | 17,66 | 430 | 466 | 468 | 469 | 469 |
| UK592_monticola_GBR | 17,95 | 597 | 605 | 605 | 606 | 606 |
| UK597_amphibius_GBR | 18,75 | 1335 | 1362 | 1362 | 1365 | 1366 |
| UK595_monticola_GBR | 19,32 | 1297 | 1337 | 1342 | 1345 | 1345 |
| UK590_monticola_GBR | 20,04 | 1023 | 1031 | 1036 | 1037 | 1038 |
| UK594_amphibius_GBR | 20,23 | 372 | 594 | 669 | 707 | 712 |
| UK588_monticola_GBR | 20,51 | 469 | 484 | 485 | 485 | 485 |
| UK584_amphibius_GBR | 21 | 71 | 153 | 184 | 201 | 211 |
| WM794_sapidus_PRT | 25,87 | 308 | 612 | 704 | 734 | 744 |
| WM710_ognevi_GEO | 27,88 | 295 | 395 | 435 | 448 | 450 |
| WM666_persicus_IRN | 28,9 | 136 | 172 | 180 | 181 | 182 |
| WM813_amphibius_BIH | 29,48 | 2 | 24 | 61 | 94 | 132 |
| WM798_amphibius_SVN | 31,45 | 86 | 338 | 467 | 559 | 613 |

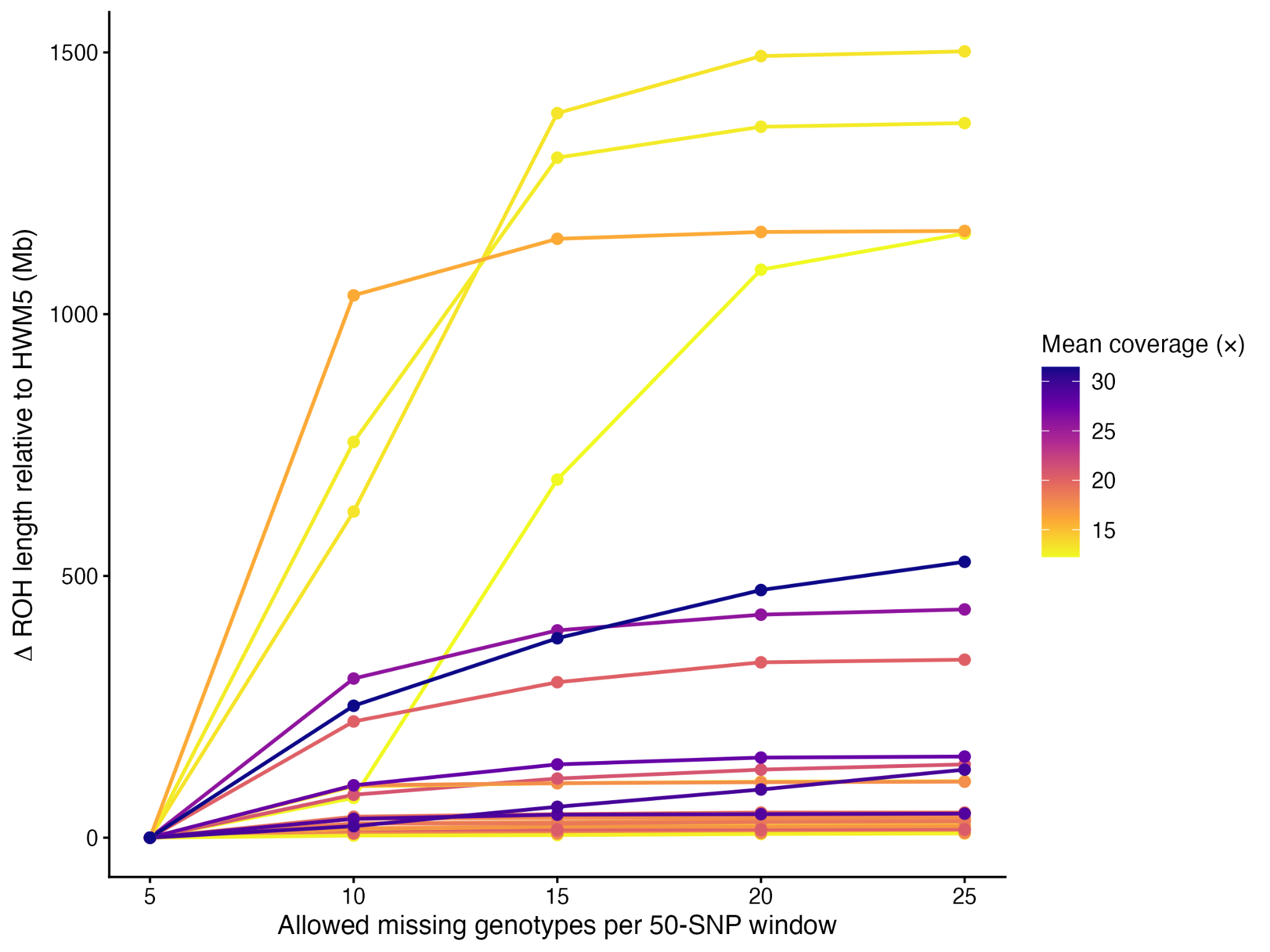

**Figure S15. Effect of the number of missing genotypes (HWM) on inferred ROH length.**

For each sample, the increase in total inferred ROH length relative to HWM5 is shown. The colours indicate mean genomic coverage.

In the case of ROHan we tested the impact of *rohmu* parameter on the recovery of ROHs as it was suggested that it may be species specific (Taylor et al. 2026). We tested values ranging from 5e−5 to 1e−3 and used 3e−4 in the final analysis (Table S10).

**Table S10.** Effect of the *rohmu* parameter on the number and total length of runs of homozygosity (ROH) inferred with ROHan reported as a fraction of genome (%) and total ROH length (Mb).

| *rohmu* | AL287 ROH (%) | AL287 ROH (Mb) | WM666 ROH (%) | WM666 ROH (Mb) |
| --- | --- | --- | --- | --- |
| 5,00E-05 | 0,0 | 0,0 | 1,6 | 33,5 |
| 1,00E-04 | 5,7 | 114,2 | 3,1 | 65,0 |
| 2,00E-04 | 12,8 | 248,8 | 4,4 | 92,5 |
| 2,50E-04 | 14,2 | 273,5 | 4,8 | 99,5 |
| **3,00E-04** | **14,9** | **282,5** | **4,8** | **104,5** |
| 3,50E-04 | 16,1 | 304,8 | 4,7 | 96,0 |
| 4,00E-04 | 16,8 | 320,8 | 4,7 | 96,0 |
| 4,50E-04 | 17,0 | 308,5 | 4,7 | 95,0 |
| 5,00E-04 | 17,8 | 319,5 | 4,7 | 94,5 |
| 1,00E-03 | 32,4 | 344,0 | 8,5 | 114,5 |

#### 2.9 Mutational load

SnpEff v.5.2. was used to estimate genetic load from three different impact classes. First, a database was built from the mArvAmp1.2 genome annotation, and used to annotate the VCF-file containing all samples >12X coverage. SnpSift v.5.0e was used to extract the relevant fields (CHROM, POS, REF, ALT, GEN [*].GT, ANN [*].IMPACT, ANN[*].EFFECT) from the annotated VCF-files, and the number of non-reference (derived) alleles in homozygous and heterozygous states was calculated for the different impact classes using custom code. The mutational load was then calculated with the formula: (counts of variants, homozygous variants counted as two)/(counts of derived alleles, homozygous alleles counted as two). The number of high, moderate, and low impact mutations per 100,000 derived alleles for each sample is reported below.

**Table S11.** Number of high, moderate, and low impact mutations per 100,000 derived alleles inferred from SnpEff.

| Sample | High impact mutations/100,000 derived alleles | Moderate impact mutations/100,000 derived alleles | Low impact mutations/100,000 derived alleles |
| --- | --- | --- | --- |
| WM666_persicus_IRN | 8,86 | 440,15 | 953,76 |
| WM710_ognevi_GEO | 8,15 | 378,52 | 809,16 |
| WM794_sapidus_PRT | 8,24 | 391,78 | 826,11 |
| WM798_amphibius_SVN | 9,17 | 428,90 | 901,45 |
| WM813_amphibius_BIH | 9,2 | 426,71 | 910,48 |
| WM833_ognevi_TUR | 9,98 | 474,25 | 1033,56 |
| AL287_italicus_ITA | 7,32 | 463,42 | 1071,84 |
| UK584_amphibius_GBR | 11,84 | 428,15 | 812,14 |
| UK588_monticola_GBR | 10,3 | 432,54 | 857,55 |
| WM818_amphibius_SVN | 11,08 | 511,0 | 1091,35 |
| UK585_monticola_GBR | 10,07 | 439,35 | 884,62 |
| UK586_monticola_GBR | 10,48 | 432,80 | 865,91 |
| UK587_monticola_GBR | 9,79 | 443,63 | 898,27 |
| UK589_monticola_GBR | 10,16 | 433,76 | 876,49 |
| UK590_monticola_GBR | 10,50 | 431,53 | 858,77 |
| UK594_amphibius_GBR | 11,55 | 427,08 | 801,42 |
| UK596_monticola_GBR | 10,23 | 429,46 | 857,64 |
| UK593_amphibius_GBR | 12,58 | 435,86 | 796,07 |
| UK595_monticola_GBR | 10,23 | 430,56 | 860,04 |
| UK597_amphibius_GBR | 11,21 | 428,31 | 818,35 |
| UK592_monticola_GBR | 10,38 | 429,26 | 862,65 |
| UK591_monticola_GBR | 9,71 | 426,20 | 860,07 |

Simonsen, Martin, Thomas Mailund, and Christian N. S. Pedersen. 2008. “Rapid Neighbour-Joining.” In *Lecture Notes in Computer Science*. Lecture Notes in Computer Science. Springer Berlin Heidelberg.

Taylor, Rebecca S., Micheline Manseau, and Paul J. Wilson. 2026. “Accurate Runs of Homozygosity Estimation from Low Coverage Genome Sequences in Non-Model Species.” *Molecular Ecology Resources* 26 (1): e70083.
